# Comparative assessment of long-read error-correction software applied to RNA-sequencing data

**DOI:** 10.1101/476622

**Authors:** Leandro Lima, Camille Marchet, Ségolène Caboche, Corinne Da Silva, Benjamin Istace, Jean-Marc Aury, Hélène Touzet, Rayan Chikhi

**Affiliations:** Univ Lyon, Université Lyon 1, CNRS, Laboratoire de Biométrie et Biologie Evolutive UMR5558 F-69622 Villeurbanne, France; EPI ERABLE - Inria Grenoble, Rhône-Alpes, France; Università di Roma “Tor Vergata”, Roma, Italy; CNRS, Université de Lille, CRIStAL UMR 9189, Lille, France; Université de Lille, CNRS, Inserm, CHU Lille, Institut Pasteur de Lille, U1019, UMR8204, Center for Infection and Immunity of Lille, Lille, France; Genoscope, Institut de biologie Francois-Jacob, Commissariat à l’Energie Atomique (CEA), Université Paris-Saclay, Evry, France; Institut Pasteur, C3BI - USR 3756, 25-28 rue du Docteur Roux, 75015 Paris, France

**Keywords:** Long reads, RNA-sequencing, Nanopore, Error correction, Benchmark

## Abstract

**Motivation:** Long-read sequencing technologies offer promising alternatives to high-throughput short read sequencing, especially in the context of RNA-sequencing. However these technologies are currently hindered by high error rates in the output data that affect analyses such as the identification of isoforms, exon boundaries, open reading frames, and the creation of gene catalogues. Due to the novelty of such data, computational methods are still actively being developed and options for the error-correction of RNA-sequencing long reads remain limited.

**Results:** In this article, we evaluate the extent to which existing long-read DNA error correction methods are capable of correcting cDNA Nanopore reads. We provide an automatic and extensive benchmark tool that not only reports classical error-correction metrics but also the effect of correction on gene families, isoform diversity, bias towards the major isoform, and splice site detection. We find that long read error-correction tools that were originally developed for DNA are also suitable for the correction of RNA-sequencing data, especially in terms of increasing base-pair accuracy. Yet investigators should be warned that the correction process perturbs gene family sizes and isoform diversity. This work provides guidelines on which (or whether) error-correction tools should be used, depending on the application type.

**Benchmarking software:** https://gitlab.com/leoisl/LR_EC_analyser

## 1. INTRODUCTION

The most commonly used technique to study transcriptomes is through RNA sequencing. As such, many tools were developed to process Illumina or short RNA-seq reads. Assembling a transcriptome from short reads is a central task for which many methods are available. When a reference genome or reference transcriptome is available, reference-based assemblers can be used (such as Cufflinks Trapnell et al. [2010], Scallop Shao and Kingsford [2017], Scripture Guttman et al. [2010], and StringTie Pertea et al. [2015]). When no references are available, *de novo* transcriptome assembly can be performed (using tools such as Oases Schulz et al. [2012], SOAPdenovo-Trans Xie et al. [2014], Trans-ABySS Robertson et al. [2010] and Trinity Grabherr et al. [2011]). Potential disadvantages of reference-based strategies include: i) the resulting assemblies might be biased towards the used reference, and true variations might be discarded in favour of known isoforms; ii) they are unsuitable for samples with a partial or missing reference genome Grabherr et al. [2011]; iii) such methods depend on correct read-to-reference alignment, a task that is complicated by splicing, sequencing errors, polyploidism, multiple read mapping, mismatches caused by genome variation, and the lack or incompleteness of many reference genomes Grabherr et al. [2011], Robertson et al. [2010]; iv) sometimes, the model being studied is sufficiently different from the reference because it comes from a different strain or line such that the mappings are not altogether reliable Schulz et al. [2012]. On the other hand, some of the shortcomings of *de novo* transcriptome assemblers are: i) low-abundance transcripts are likely to not be fully assembled Haas and Zody [2010]; ii) reconstruction heuristics are usually employed, which may lead to missing alternative transcripts, and highly similar transcripts are likely to be assembled into a single transcript Martin and Wang [2011]; (iii) homologous or repetitive regions may result in incomplete assemblies Fu et al. [2018]; (iv) accuracy of transcript assembly is called into question when a gene exhibits complex isoform expression Fu et al. [2018].

Recent advances in long-read sequencing technology have enabled longer, up to full-length sequencing of RNA molecules. This new approach has the potential to eliminate the need for transcriptome assembly, and thus also eliminate from transcriptome analysis pipelines all the biases caused by the assembly step. Long read sequencing can be done using either cDNA-based or direct RNA protocols from Oxford Nanopore (referred to as *ONT* or *Nanopore*) and Pacific Biosciences (*PacBio*). The Iso-Seq protocol from PacBio consists in a size selection step, sequencing of cDNAs, and finally a set of computational steps that produce sequences of full-length transcripts. ONT has three different experimental protocols for sequencing RNA molecules: cDNA transformation with amplification, direct cDNA (with or without amplification), and direct RNA.

Long-read sequencing is increasingly used in transcriptome studies, not just to prevent problems caused by short-read transcriptome assembly, but also for several of the following reasons. Mainly, long reads can better describe exon/intron combinations Sedlazeck et al. [2018]. The Iso-seq protocol has been used for isoform identification, including transcripts identification Wang et al. [2016], de novo isoform discovery Li et al. [2017] and fusion transcript detection Weirather et al. [2015]. Nanopore has recently been used for isoform identification Byrne et al. [2017] and quantification Oikonomopoulos et al. [2016].

The sequencing throughput of long-read technologies is significantly increasing over the years. It is now conceivable to sequence a full eukaryote transcriptome using either only long reads, or a combination of high-coverage long and short (Illumina) reads. Unlike the Iso-Seq protocol that requires extensive *in silico* processing prior to primary analysis Sahlin et al. [2018], raw Nanopore reads can in principle be readily analyzed. Direct RNA reads also permit the analysis of base modifications Workman et al. [2018], unlike all other cDNA-based sequencing technologies. There also exist circular sequencing techniques for Nanopore such as INC-Seq Li et al. [2016] which aim at reducing error rates, at the expense of a special library preparation. With raw long reads, it is up to the primary analysis software (typically a mapping algorithm) to deal with sequences that have significant per-base error rate, currently around 13% Weirather et al. [2017].

In principle, a high error rate in the data complicates the analysis of transcriptomes especially for the accurate detection of exon boundaries, or the quantification of similar isoforms and paralogous genes. Reads need to be aligned unambiguously and with high base-pair accuracy to either a reference genome or transcriptome. Indels (i.e. insertions/deletions) are the main type of errors produced by long-read technologies, and they confuse aligners more than substitution errors Sović et al. [2016]. Many methods have been developed to correct errors in RNA-seq reads, mainly in the short-read era Tong et al. [2016], Song and Florea [2015]. They no longer apply to long reads because they were developed to deal with low error rates, and principally substitutions. However, a new set of methods have been proposed to correct genomic long reads. There exist two types of long-read error-correction algorithms, those using information from long reads only (*self* or *non-hybrid* correction), and those using short reads to correct long reads (*hybrid* correction). In this article, we will report on the extent to which state-of-the-art tools enable to correct long noisy RNA-seq reads produced by Nanopore sequencers.

Several tools exist for error-correcting long reads, including ONT reads. Even if the error profiles of Nanopore and PacBio reads are different, the error rate is quite similar and it is reasonable to expect that tools originally designed for PacBio data to also perform well on recent Nanopore data. There is, to the best of our knowledge, very little prior work that specifically addresses error-correction of RNA-seq long reads. Notable exceptions include: a) LSC Au et al. [2012], which is designed to error correct PacBio RNA-seq long reads using Illumina RNA-seq short reads; b) PBcR Koren et al. [2012] and c) HALC Bao and Lan [2017], which are mainly designed for genomes but are also evaluated on transcriptomic data. Here we will take the standpoint of evaluating long-read error-correction tools on RNA-seq data, most of which were designed to process DNA sequencing data only.

We evaluate the following DNA hybrid correction tools: HALC Bao and Lan [2017], LoRDEC Salmela and Rivals [2014], NaS Madoui et al. [2015], PBcR Koren et al. [2012], proovread Hackl et al. [2014]; and the following DNA self-correction tools: Canu Koren et al. [2017], daccord Tischler and Myers [2017], LoRMA Salmela et al. [2016], MECAT Xiao et al. [2017], pbdagcon Chin et al. [2013]. We also evaluate an additional hybrid tool, LSC Au et al. [2012], the only one specifically designed to error correct (PacBio) RNA-seq long reads.

A majority of hybrid correction methods employ mapping strategies to place short fragments on long reads and correct long read regions using the related short read sequences. But some of them rely on graphs to create a consensus that is used for correction. These graphs are either k-mer graphs (de Bruijn graphs), or nucleotide graphs resulting from multiple alignments of sequences (partial order alignment). For self-correction methods, strategies using the aforementioned graphs are the most common. We have also considered evaluating nanocorrect Loman et al. [2015], nanopolish Loman et al. [2015], Falcon sense Chin et al. [2016], and LSCPlus Hu et al. [2016], but some tools were deprecated, not suitable for read correction, or unavailable. Our detailed justifications can be found in Section S1.12 of the Supplementary Material. We have selected what we believe is a representative set of tools but there also exist other tools that were not considered in this study, e.g. HG-Color Morisse et al. [2018], HECIL Choudhury et al. [2018], MIRCA Kchouk and Elloumi [2016], Jabba Miclotte et al. [2016], nanocorr Goodwin et al. [2015], and Racon Vaser et al. [2017].

Other works have evaluated error correction tools in the context of DNA sequencing. LRCstats La et al. [2017], and more recently ELECTOR Marchet et al. [2019], provide automated evaluations of genomic long read correction using a simulated framework. A technical report from Bouri and Lavenier [2017] provides an extensive evaluation of PacBio/Nanopore error-correction tools, in the context of de novo assembly. This analysis is completed with more recent results in Fu et al. [2019] on hybrid correction methods. Perhaps the closest work to ours is the AlignQC software Weirather et al. [2017], which provides a set of metrics for the evaluation of RNA-sequencing long-read dataset quality. In Weirather et al. [2017] a comparison is provided between Nanopore and PacBio RNA-sequencing datasets in terms of error patterns, isoform identification and quantification. While Weirather et al. [2017] did not compare error-correction tools, we will use and extend AlignQC metrics for that purpose.

In this article, we will focus on the qualitative and quantitative measurements of error-corrected long reads, with transcriptomic features in mind. First we examine basic metrics of error-correction, e.g. mean length, base accuracy, homopolymers errors, and performance (running time, memory) of the tools. Then we ask several questions that are specific to transcriptome applications: (i) how is the number of detected genes, and more precisely the number of genes within a gene family, impacted by read error correction? (ii) can error correction significantly change the number of reads mapping to genes or transcripts, possibly affecting downstream analysis based on these metrics? (iii) do error-correction tools perturb isoform diversity, e.g. by having a correction bias towards the major isoform? (iv) what is the impact of error correction on identifying splice sites? To answer these questions, we provide an automatic framework (LC EC analyser, see Methods) for the evaluation of transcriptomic error-correction methods, that we apply to eleven different error-correction tools.

## 2. RESULTS

### 2.1 Error-correction tools

Table 1 presents the main characteristics of the hybrid and non-hybrid error-correction tools that were considered in this study. For the sake of reproducibility, in the Supplementary Material Section S1 are described all the versions, dependencies, and parameters. Note that these error-correction tools were all tailored for DNA-seq data except for LSC.

**Table 1.** Main characteristics of the error correction tools considered in this study

(A) Hybrid tools
|  | HALC | LoRDEC | LSC | NaS | PBcR | Proovread |
| --- | --- | --- | --- | --- | --- | --- |
| Reference | Bao and Lan [2017] | Salmela and Rivals [2014] | Au et al. [2012] | Madoui et al. [2015] | Koren et al. [2012] | Hackl et al. [2014] |
| Context | RNA or DNA | DNA | RNA | DNA | RNA or DNA | DNA |
| Technology | PacBio | PacBio or ONT | PacBio | ONT | PacBio or ONT | PacBio |
| Main algorithmic idea | Aligns short reads to long reads. HALC implements a strategy to desambiguify multiple alignments instead of avoiding them. It uses a short reads contig graph in order to select the best set of contigs to correct a long read region. | Construction of short read de Bruijn graph (DBG), path search between k-mers in long reads. Regions between k-mers are corrected with the optimal path. | Operate a homopolymer compression of short and long reads to increase alignment recall. Recruits short reads on long reads sequences. Corrects errors from short reads sequences and uses homopolymers from short reads to replace those in long reads. | Long reads are used as templates to recruit short (seed) reads through alignment. The set of seeds is extended by searching, using shared k-mers, for similar reads in the initial read set. A micro-assembly of the reads is performed. Resulting contigs are aligned back to the input long reads, and a path in the contig graph is used as the corrected read. | Alignment of short reads to long reads, and multi-alignment of the short reads recruited to one region. A consensus is derived from the multiple alignment. | Alignment of short reads to long reads and consensus. Uses a specific scoring system to adapt the mapping to the high error rates. |

(B) Non-hybrid tools
|  | Canu | daccord | LoRMA | MECAT | pbdagcon |
| --- | --- | --- | --- | --- | --- |
| Reference | Koren et al. [2017] | Tischler and Myers [2017] | Salmela et al. [2016] | Xiao et al. [2017] | Chin et al. [2013] |
| Context | DNA | DNA | DNA | DNA | DNA |
| Technology | PacBio or ONT | PacBio | PacBio or ONT | PacBio or ONT | PacBio |
| Main algorithmic idea | First, All-versus-all read overlap and alignment filtering. Directed acyclic graph is built from the alignments, that produce quasi-multiple alignments. Highest-weight path search in these graph yield consensus sequences that are used for correction. | Reads are compared pairwise to obtain alignment piles. Several overlapping windows of hundreds of nucleotides are derived from these piles, on which micro-assembly (DBG with very small k-mers) is performed. A heaviest path in each DBG is heuristically selected as the consensus to correct the given window. | Path search in DBG built from long reads. Multi-iterations over k-mer size for graph construction. The same framework than LoRDEC is used to correct read regions. | k-mer based read matching, pairwise alignment between matched reads, alignment-based consensus calling on 'easy' regions, local consensus calling (partial order graph) otherwise. | Align long reads to "backbone" sequences, correction by iterative directed acyclic graph consensus calling from the multiple sequence alignments. |

The output of each error correction method can be classified into on of the four following types: full-length, trimmed, split, and micro-assembly. Usually, due to methodological reasons, extremities of long reads are harder to correct. As an example, hybrid correctors based on mapping short to long reads, and calling a consensus from the mapping, have difficulties aligning short reads to the extremities of long reads. As such, some methods output *trimmed error-corrected reads*, i.e. error-corrected reads such that their uncorrected ends are removed. Examples of methods producing this type of output considered in this study are HALC, LoRDEC, LSC, proovread, daccord, and pbdagcon. Sometimes, internal parts of long reads can also be hard to correct, due to a lack of coverage of short reads, or a drop of sequencing quality, or due to mapping issues. Some algorithms thus output *split error-corrected reads*, splitting one long read into several well-corrected fragments, such as HALC, LoRDEC, PBcR, and LoRMA. Finally, some tools decide to not trim nor split the original reads, outputting *full-length error-corrected reads*. Examples include HALC, LoRDEC, LSC, proovread, canu, daccord, MECAT, and pbdagcon. NaS does not fit the previous three categories, as it uses a micro-assembly strategy, instead of a classical polishing of the consensus, in which the long read is used as a template to recruit Illumina reads and, by performing a local assembly, build a high-quality synthetic read. As can be noted, some tools produce more than one type of output, sometimes three types. In the following sections, we add the suffixes *(t)* and *(s)* to a tool name to denote its trimmed and split outputs, respectively. We further add the suffix *(µ)* to NaS, as a reminder that it is based on a micro-assembly strategy. Outputs that have no suffixes are considered full-length corrections. For example, HALC denotes the HALC full-length error-corrected reads, HALC(t), the HALC trimmed output, and HALC(s), the HALC split output. As we will see, there is no type of output that outperforms all the others in all metrics. Choosing the appropriate type of output is heavily dependent on the application.

### 2.2 Evaluation datasets

Our main evaluation dataset consists of a single 1D Nanopore run using the cDNA preparation kit of RNA material taken from a mouse brain, containing 740,776 long reads. An additional Illumina dataset containing 58 million paired-end 151 bp reads was generated on the same sample but using a different cDNA protocol. For more details on the sequencing protocol, see Section 4. The Nanopore and Illumina reads from the mouse RNA sample are available in the ENA repository under the following study: PRJEB25574. In this paper, we provide a detailed analysis of this dataset, from Section 2.3 to Section 2.11.

In order to obtain a more comprehensive understanding of the evaluated tools, we further analysed the correction of the methods on one human Nanopore direct RNA sequencing data from the Nanopore-WGS-Consortium (dataset from centre Bham, run#1, sample type RNA, kit SQK-RNA001, pore R9.4, available at https://github.com/nanopore-wgs-consortium/NA12878/blob/master/nanopore-human-transcriptome/fastq_fast5_bulk.md). We concatenated the fail and pass RNA-direct reads from the aforementioned dataset, obtaining 894,289 reads. Further, to correctly run all tools, we transformed bases U into T.

### 2.3 Error-correction improves base accuracy and splits, trims, or entirely removes reads

Table 2 shows an evaluation of error-correction based on AlignQC results, for the hybrid and non-hybrid tools. The error rate is 13.72% in raw reads, 0.33-5.45% for reads corrected using hybrid methods and 2.91-6.43% with self-correctors. Notably, the hybrid tools NaS(*µ*), Proovread(t), and HALC(s) output micro-assembled, trimmed and split error-corrected reads, respectively, with the lowest error rates (¡0.5%). We observe that HALC produced the full-length error-corrected reads with the lowest error-rate (1.85%), but that is still significantly higher than the error-rate of the three aforementioned methods. This is expected, as micro-assembling, trimming or splitting reads usually do not retain badly corrected regions of the reads, lowering the error rate. LoRMA(s), which is the only split self-correction tool, was the one that decreased the error-rate the most among non-hybrid tools, but still just managed to reach 2.91%, one order of magnitude higher than the best hybrid correctors. If we look at non-split outputs among the self-correctors, MECAT and daccord(t) obtained the lowest error rates for full-length and trimmed error-corrected reads, respectively, but still presenting an error-rate higher that 4%. It is not a surprise that the best error correctors are hybrid when looking at the error rates, given their usage of additional high-quality Illumina reads. As expected, trimming and splitting error-corrected reads reduces a lot the error-rate, e.g. LoRDEC reduced the error rate from 4.5% to 3.73% by trimming, and to 1.59% by splitting. As such, the split output consistently outperformed trimmed and full-length outputs, regarding the error-rate. A detailed error-rate analysis will be carried in Section 2.5.

**Table 2.** Statistics of error correction tools on the 1D run RNA-seq dataset. To facilitate the readability of this table and the next ones, we highlighted values that we deemed satisfactory in green colour, borderline in brown, and unsatisfactory in red, noting that such classification is somewhat arbitrary.

| | Raw | HALC | HALC(t) | HALC(s) | LoRDEC | LoRDEC(t) | LoRDEC(s) | LSC | LSC(t) | NaS($\mu$) | PBcR(s) | Proovread | Proovread(t) |
| --- | --- | --- | --- | --- | --- | --- | --- | --- | --- | --- | --- | --- | --- |
| nb reads | 741k | 741k | 709k | 914k | 741k | 677k | 1388k | 619k | 619k | 619k | 1321k | 738k | 626k |
| mapped reads | 83.5% | 88.1% | 95.6% | 98.8% | 85.5% | 95.5% | 97.5% | 97.1% | 97.6% | 98.7% | 99.2% | 85.5% | 98.9% |
| mean length | 2011 | 2174 | 1926 | 1378 | 2097 | 1953 | 816 | 2212 | 1901 | 1931 | 776 | 2117 | 1796 |
| nb bases | 1313M | 1469M | 1334M | 1245M | 1394M | 1289M | 1106M | 1332M | 1151M | 1179M | 1015M | 1400M | 1112M |
| mapped bases<sup>a</sup> | 89.0% | 90.3% | 96.6% | 99.2% | 90.6% | 95.9% | 99.1% | 90.9% | 97.7% | 97.5% | 99.2% | 92.4% | 99.5% |
| error rate<sup>b</sup> | 13.72% | 1.85% | 1.32% | 0.44% | 4.5% | 3.73% | 1.59% | 5.45% | 4.36% | 0.38% | 0.68% | 2.65% | 0.33% |
| nb detected genes | 16.8k | 17.0k | 16.8k | 16.6k | 16.8k | 16.6k | 16.5k | 16.5k | 16.2k | 15.0k | 15.6k | 16.6k | 14.6k |

(B) Non-hybrid tools
|  | Raw | Canu | daccord | daccord(t) | LoRMA(s) | MECAT | pbdagcon | pbdagcon(t) |
| --- | --- | --- | --- | --- | --- | --- | --- | --- |
| nb reads | 741k | 519k | 675k | 840k | 1540k | 495k | 778k | 775k |
| mapped reads | 83.5% | 99.1% | 92.5% | 94.0% | 99.4% | 99.4% | 98.2% | 97.9% |
| mean length | 2011 | 2192 | 2102 | 1476 | 497 | 1992 | 1473 | 1484 |
| nb of bases | 1313M | 1125M | 1350M | 1212M | 760M | 980M | 1136M | 1141M |
| mapped bases<sup>a</sup> | 89.0% | 92.0% | 92.5% | 94.7% | 99.2% | 96.9% | 97.0% | 96.7% |
| error rate<sup>b</sup> | 13.72% | 6.43% | 5.2% | 4.12% | 2.91% | 4.57% | 5.65% | 5.71% |
| nb detected genes | 16.8k | 12.4k | 15.5k | 13.9k | 6.8k | 10.4k | 13.2k | 13.2k |

<sup>a</sup>As reported by AlignQC. Percentage of bases aligned among mapped reads, taken by counting the M parts of CIGAR strings in the BAM file. Bases in unmapped reads are not counted.
<sup>b</sup>As reported by AlignQC, using a sample of 1 million bases from aligned reads segments.

In terms of throughput after the correction step, tools that do not trim nor split reads tend to return a number of reads similar to that of the uncorrected (raw) reads. Notably, HALC and LoRDEC returned exactly the same number of reads, and Proovread returned just 3k less reads. On the other hand, Canu and MECAT decreased a lot the number of output reads, probably due to post-filtering procedures. Moreover, many of full-length outputs (HALC, LoRDEC, LSC, Proovread, and daccord) increased the mean length of the raw reads while also increasing the number of output bases, showing that they tend to further extend the information contained in the long reads.

Trimming almost always decreased the number of output reads, like in HALC(t), LoRDEC(t), Proovread(t), and pbdagcon(t), probably due to post-filtering procedures. However, in LSC(t), trimming has no effect on the number of reads, and in daccord(t), trimming actually increased the number of reads. In half of the trimmed outputs (HALC(t), LoRDEC(t), and LSC(t)), the mean length of the reads was usually preserved, decreasing only by around 100bps. However, in the other half (Proovread(t), daccord(t), and pbdagcon(t)), read lengths were, on average, reduced by 200-500bp.

Splitting reads significantly increased the number of output reads, as expected. LoRDEC(s), PBcR(s), and LoRMA(s) tend to split reads into two or more shorter reads during the correction step, as they return ∼2× more reads after correction that are also shorter (mean length of respectively 816bp, 776bp and 497bp, versus 2011bp in raw reads) and overall have significantly less bases in total (loss of respectively 207Mbp, 298Mbp and 553Mbp). HALC(s), on the other hand, managed to increase the number of reads only by 23%, with no significant loss of bases, but still with a significant reduction on the mean length (1378bp). We observe that NaS(*µ*), based on micro-assembly, obtained a mean length similar to the trimmed outputs (HALC(t), LoRDEC(t), and LSC(t)). This suggests that NaS(*µ*) has trouble either getting seed short reads or recruiting short reads mapping to the ends of long reads, or assembling the reads mapped to the ends.

These observations indicate that care should be taken when considering which type of output should be used. For example, all split and half of the trimmed outputs should not be used in applications trying to describe the full transcript structure, or distant exons coupling, as the long read connectivity is lost in many cases in these types of outputs.

Overall, no correction tool outperforms all the others across the metrics analysed in this section. However, hybrid correctors are systematically better than self-correctors at decreasing the error-rate (and preserving the transcriptome diversity, as we will discuss in the next Section). Trimming and splitting usually increase the read accuracy (and also mapping rate, as we see next), but decrease the total amount of bases in the read set and the mean read length, which can lead to loss of long-range information that was present in the raw reads.

### 2.4 Error-correction facilitates mapping yet generally lowers the number of detected genes

Apart from HALC, LoRDEC, Proovread, and daccord, for which only 85-92% of reads were mapped, corrected reads from all the other tools were mapped at a rate of 94-99%, showing a significant improvement over raw reads (mapping rate of 83.5%). We observe that these four tools with the lowest percentages of mapped reads had high mean read length, indicating that trimming, splitting or discarding reads seems necessary in order to obtain shorter but overall less error-prone reads. In general in all tools (except pbdagcon), trimming and splitting increased the proportion of mapped reads and bases, sometimes significantly (e.g. Proovread). However some tools which do not offer the option to trim or split reads, such as Canu and MECAT, showed very high mapping rate with high mean read length and error-rate. This is related to their aggressive post-filtering measure, which removed a significant portion of the reads (29-33%).

On verifying if error-correctors are able to preserve transcriptome diversity, we can see a striking difference between hybrid and self-correctors: in general, hybrid correctors present a far higher number of detected genes than the self ones. Interestingly, HALC was able to even increase the number of detected genes by 221 with regard to the raw reads, indicating that some genes were maybe not detected before due to imperfect mapping caused by the high error rate. We also found that 72 genes were detected in the raw reads but not in any of the error-corrected outputs. Furthermore, 354 genes are absent from the results of nearly all correction methods (≥16 out of 19).

Overall, all hybrid tools presented a satisfactory amount of detected genes, except for NaS(*µ*), PBcR(s) and Proovread (t), while self-correctors did not present any satisfactory results, with Canu, LoRMA(s) and MECAT reducing by 35%-59% the number of detected genes reported in raw reads. We can also note that trimming and splitting systematically resulted in a loss of the sensitivity to detect new genes. Moreover, except for HALC(s), tools with very high percentage of mapped reads (NaS(*µ*), PBcR(s), Proovread(t), Canu, LoRMA(s), MECAT, pbdagcon, pbdagcon(t)) had the largest losses in number of detected genes, hinting that error correction can reduce gene diversity in favor of lower error-rate, and/or that clusters of similar genes (e.g. paralogous) are corrected towards a single gene. Therefore, if preserving the transcriptome diversity is required for the downstream application, self-correctors should be avoided altogether, along with some hybrid correctors (NaS(*µ*), PBcR(s), and Proovread(t)).

### 2.5 Detailed error-rate analysis

The high error-rate of transcriptomic long reads significantly complicates their primary analysis Križanović et al. [2018]. While Section 2.3 presented a general per-base error rate, this section breaks down sequencing errors into several types and examines how each error-correction tool deals with them. A general, and expected, trend that we find in all tools and in all types of errors is that trimming and splitting the reads result in less substitutions, deletions and insertion errors. We will therefore focus in other aspects in this analysis. The data presented here is a compilation of AlignQC results. Note that AlignQC computed the following metrics only on reads that could be aligned, thus unaligned reads are not counted, yet they may possibly be the most erroneous ones. AlignQC also subsampled aligned reads to around 1 million bases to calculate the presented values.

#### 2.5.1 Deletions are the most problematic sequencing errors

Table 3 shows the error rate in the raw reads and in the corrected reads for each tool. In raw reads, deletions are the most prevalent type of errors (7.41% of bases), closely followed by subsitutions (5.11%), then insertions (1.2%). LoRDEC, LSC and LSC(t) are the least capable of correcting mismatches (¿2% of them remaining), even though they are all hybrid tools. For LoRDEC, we were able to verify that this is related to the large amount of uncorrected reads in its output (90k totally uncorrected reads out of 741k - 12%), as computed by exactly matching raw reads to its corrected output. For LSC and LSC(t), we were unable to pinpoint a reason. The majority of other hybrid tools (HALC, HALC(t), HALC(s), NaS(*µ*), PBcR(s), Proovread, Proovread(t)) result in less than 1% of substitution errors. Surprisingly, the non-hybrid tools also presented very low mismatches rates: all of them showed rates lower than 1%, except for Canu (1.33%) and daccord (1.1%). This suggests that the rate of systematic substitution errors in ONT data is low, as self-correctors were able to achieve results comparable to the hybrid ones, even without access to Illumina reads. Still, the three best performing tools were all hybrid (NaS(*µ*), PBcR(s), and Proovread(t)), which should therefore be preferred for applications that require very low mismatch rates.

**Table 3.**
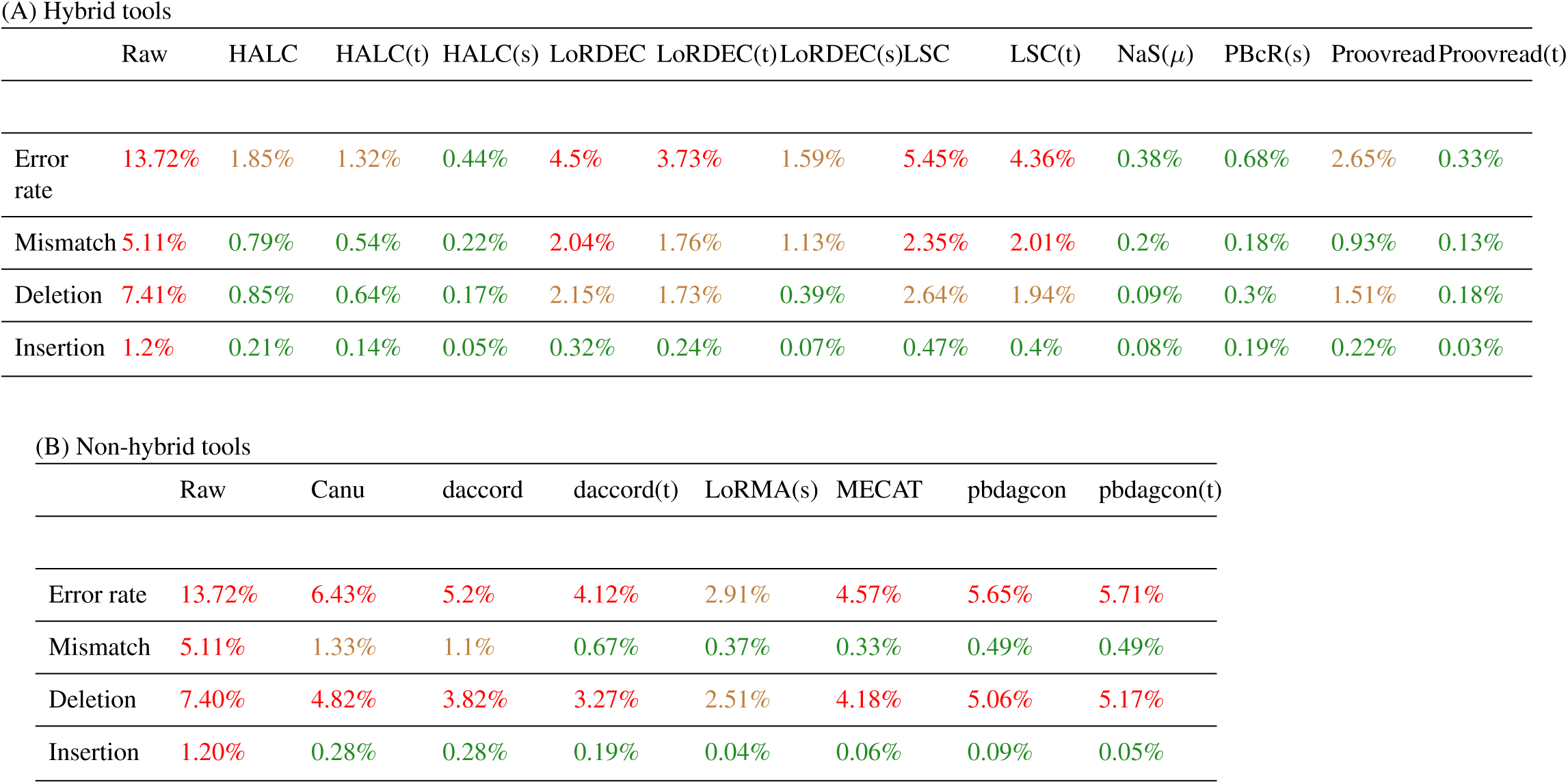
Error rate in the raw reads and in the corrected reads for each tool, on the 1D run RNA-seq dataset, computed from 1M random aligned bases.

| (A) Hybrid tools |  |  |  |  |  |  |  |  |  |  |  |  |  |
| --- | --- | --- | --- | --- | --- | --- | --- | --- | --- | --- | --- | --- | --- |
| | Raw | HALC | HALC(t) | HALC(s) | LoRDEC | LoRDEC(t) | LoRDEC(s) | LSC | LSC(t) | NaS($\mu$) | PBcR(s) | Proovread | Proovread(t) |
| Error rate | 13.72% | 1.85% | 1.32% | 0.44% | 4.5% | 3.73% | 1.59% | 5.45% | 4.36% | 0.38% | 0.68% | 2.65% | 0.33% |
| Mismatch | 5.11% | 0.79% | 0.54% | 0.22% | 2.04% | 1.76% | 1.13% | 2.35% | 2.01% | 0.2% | 0.18% | 0.93% | 0.13% |
| Deletion | 7.41% | 0.85% | 0.64% | 0.17% | 2.15% | 1.73% | 0.39% | 2.64% | 1.94% | 0.09% | 0.3% | 1.51% | 0.18% |
| Insertion | 1.2% | 0.21% | 0.14% | 0.05% | 0.32% | 0.24% | 0.07% | 0.47% | 0.4% | 0.08% | 0.19% | 0.22% | 0.03% |

| (B) Non-hybrid tools |  |  |  |  |  |  |  |  |
| --- | --- | --- | --- | --- | --- | --- | --- | --- |
|  | Raw | Canu | daccord | daccord(t) | LoRMA(s) | MECAT | pbdagcon | pbdagcon(t) |
| Error rate | 13.72% | 6.43% | 5.2% | 4.12% | 2.91% | 4.57% | 5.65% | 5.71% |
| Mismatch | 5.11% | 1.33% | 1.1% | 0.67% | 0.37% | 0.33% | 0.49% | 0.49% |
| Deletion | 7.40% | 4.82% | 3.82% | 3.27% | 2.51% | 4.18% | 5.06% | 5.17% |
| Insertion | 1.20% | 0.28% | 0.28% | 0.19% | 0.04% | 0.06% | 0.09% | 0.05% |

The contrast between self and hybrid tools is more visible on deletion errors. In general, all hybrid tools outperformed the non-hybrid ones (the only exception is LSC (2.64%), with higher deletion error rate than LoRMA(s) (2.51%)). Although in the hybrid ones, LoRDEC (2.15%), LSC (2.64%), LSC(t) (1.94%) and Proovread (1.51%) still showed moderate rates of deletions, all the other seven corrected outputs were able to lower the deletion error rate from 7.4% to less than 1%. Notably, HALC(s) and Proovread(t) to less than 0.2%, and NaS(*µ*) to less than 0.1%. All non-hybrid tools presented a high rate (3% or more) of deletion errors, except LoRMA(s) (2.51%). This comparison suggests that ONT reads exhibit systematic deletions, that cannot be well corrected without the help of Illumina data. The contribution of homopolymer errors will be specifically analyzed in Section 2.5.2. Considering insertion errors, all tools performed equally well. It is worth noting that several hybrid (HALC(s), LoRDEC(s), NaS(*µ*), and Proovread(t)) and non-hybrid tools (LoRMA(s), MECAT, pbdagcon, and pbdagcon(t)) achieved sub-0.1% insertion rate errors.

Overall, hybrid tools outperformed non-hybrid ones in terms of error-rate reduction. However, the similar results obtained by both types of tools when correcting mismatches and insertions, and the contrast in correcting deletions, seem to indicate that the main advantage of hybrid correctors over self-correctors is the removal of systematic errors using Illumina data.

#### 2.5.2 Homopolymer insertions are overall better corrected than deletion

In this section we further analyze homopolymers indels, *i.e.* insertion or deletion errors consisting of a stretch of the same nucleotide. Table 4 shows that homopolymer deletions are an order of magnitude more abundant in raw reads than homopolymer insertions. It is worth noting that, by comparing the values for the raw reads in Tables 3 and 4, homopolymers are involved in 40% of all deletions and 31% of all insertions.

**Table 4.** Homopolymer error rate in the raw reads and in the corrected reads for each tool, on the 1D run RNA-seq dataset, computed from 1M random aligned bases.

| (A) Hybrid tools |  |  |  |  |  |  |  |  |  |  |  |  |  |
| --- | --- | --- | --- | --- | --- | --- | --- | --- | --- | --- | --- | --- | --- |
| | Raw | HALC | HALC(t) | HALC(s) | LoRDEC | LoRDEC(t) | LoRDEC(s) | LSC | LSC(t) | NaS($\mu$) | PBcR(s) | Proovread | Proovread(t) |
| Homop. deletion | 2.96% | 0.28% | 0.19% | 0.03% | 0.77% | 0.63% | 0.19% | 0.62% | 0.42% | 0.02% | 0.1% | 0.46% | 0.04% |
| Homop. insertion | 0.38% | 0.05% | 0.03% | 0.01% | 0.09% | 0.07% | 0.02% | 0.11% | 0.09% | 0.01% | 0.02% | 0.06% | 0.01% |

| (B) Non-hybrid tools |  |  |  |  |  |  |  |  |
| --- | --- | --- | --- | --- | --- | --- | --- | --- |
|  | Raw | Canu | daccord | daccord(t) | LoRMA(s) | MECAT | pbdagcon | pbdagcon(t) |
| Homop. deletion | 2.96% | 2.46% | 2.14% | 2.05% | 1.82% | 2.09% | 2.26% | 2.27% |
| Homop. insertion | 0.38% | 0.08% | 0.06% | 0.03% | 0.01% | 0.01% | 0.02% | 0.01% |

A closer look at Table 4 reveals that hybrid error correctors outperform non-hybrid ones, as expected, mainly as homopolymer indels are likely systematic errors in ONT sequencing. Hybrid correctors correct them using Illumina reads that do not contain such biases. Moreover, all tools performed well on correcting homopolymer insertions, reducing the rate from 0.38% to less than 0.11%. In particular, the hybrid tools HALC(s), NaS(*µ*) and Prooovread(t), as well as the non-hybrid ones LoRMA(s), MECAT and pbdagcon(t) reached 0.01% homopolymer insertion error rate. Regarding homopolymer deletions, the majority of hybrid tools returned less than 0.5% of them, except LoRDEC (0.77%), LoRDEC(t) (0.63%), and LSC (0.62%). Notably, HALC(s), NaS(*µ*), and Proovread(t) presented less than 0.05% of homopolymer deletion error rate. Non-hybrid tools performed more pooly, returning 1.8-2.4% of homopolymers deletion errors – a small improvement over the raw reads.

HALC(s), NaS(*µ*) and Proovread(t) showed the best reduction of homopolymers indels. It is also worth noting that hybrid correctors are able to correct homopolymer deletions even better than non-homopolymer deletions. For instance the ratio of homopolymer deletions over all deletions is 39.9% in raw reads, and decreases for all hybrid correctors, except LoRDEC(s), dropping to 17.6% for HALC(s), and 22.2% for NaS(*µ*) and Proovread(t), but increases to at least 43.9% (pbdagcon(t)) up to 72.5% (LoRMA(s)) in non-hybrid tools (see Supplementary Material Section S2).

### 2.6 Error-correction perturbs the number of reads mapping to the genes and transcripts

Downstream RNA-sequencing analyses typically rely on the number of reads mapping to each gene and transcript for quantification, differential expression analysis, etc. In the rest of the paper, we define the **coverage** of a gene or a transcript as the number of reads mapping to it. For short we will refer to those coverages as *C*_*G*_ and *C*_*T*_, respectively. In this section we investigate if the process of error correction can perturb *C*_*G*_ and *C*_*T*_, which in turn would affect downstream analysis. Note that error correction could potentially slightly increase coverage, as uncorrected reads that were unmapped can become mappable after correction. Figure 1 shows the *C*_*G*_ before and after correction for each tool. We can note, as expected, that splitting a long read into several well-corrected fragments generates multiple counts, skewing up the observed coverages (see HALC(s), LoRDEC(s), PBcR(s), LoRMA(s) in Figure 1). Therefore, users are recommended to not use this type of output for gene coverage estimation. Further, apart from the split outputs, all the other tools presented good correlation and the expected slight increase in *C*_*G*_ due to better mapping, except for MECAT, which presented the lowest correlation and a significant drop in *C*_*G*_. All tools systematically presented a similar trend and lower correlation values on *C*_*T*_ (see Supplementary Material Section S3), in comparison to *C*_*G*_. This is expected, as it is harder for a tool to correct a read into its true isoform than into its true gene. The behaviour of the tools in the isoform level are in coherence with their behaviour in the gene level (*C*_*G*_): split outputs inflate *C*_*T*_; MECAT deflates it; and all the others present a slight increase.

**Fig. 1.**
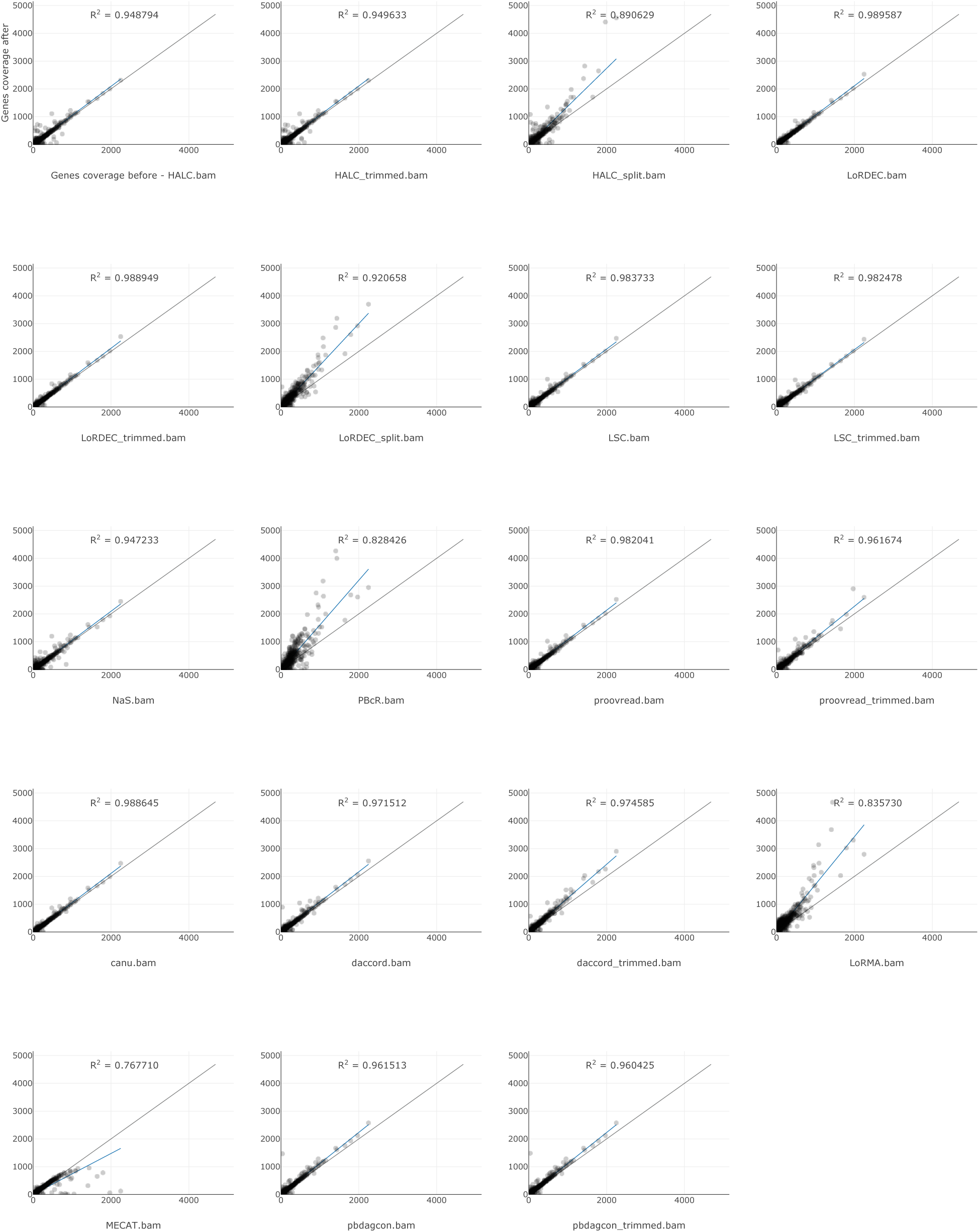
Number of reads mapping to genes (*C*_*G*_) before and after correction for each tool. The genes taken into account here were expressed in either the raw dataset or after the correction by the given tool.

### 2.7 Error-correction perturbs gene family sizes

Table 2 indicates that error correction generally results in a lower number of detected genes. In this section we explore the impact of error-correction on paralogous genes. By paralogous **gene family**, we denote a set of paralogs computed from Ensembl (see Section 4.3). Figure 2 represents the changes in sizes of paralogous gene families before and after correction for each tool, in terms of number of genes expressed within a given family. Overall, error-correctors do not strictly preserve the sizes of gene families. Correction more often shrinks families of paralogous genes than it expands them, likely due to erroneous correction in locations that are different between paralogs. In summary, 36-87% of gene families are kept of the same size by correctors, 1-17% are expanded and 6-61% are shrunk. Supplementary Material Figure S2 shows the magnitude of expansion/shrinkage for each gene family.

**Fig. 2.**
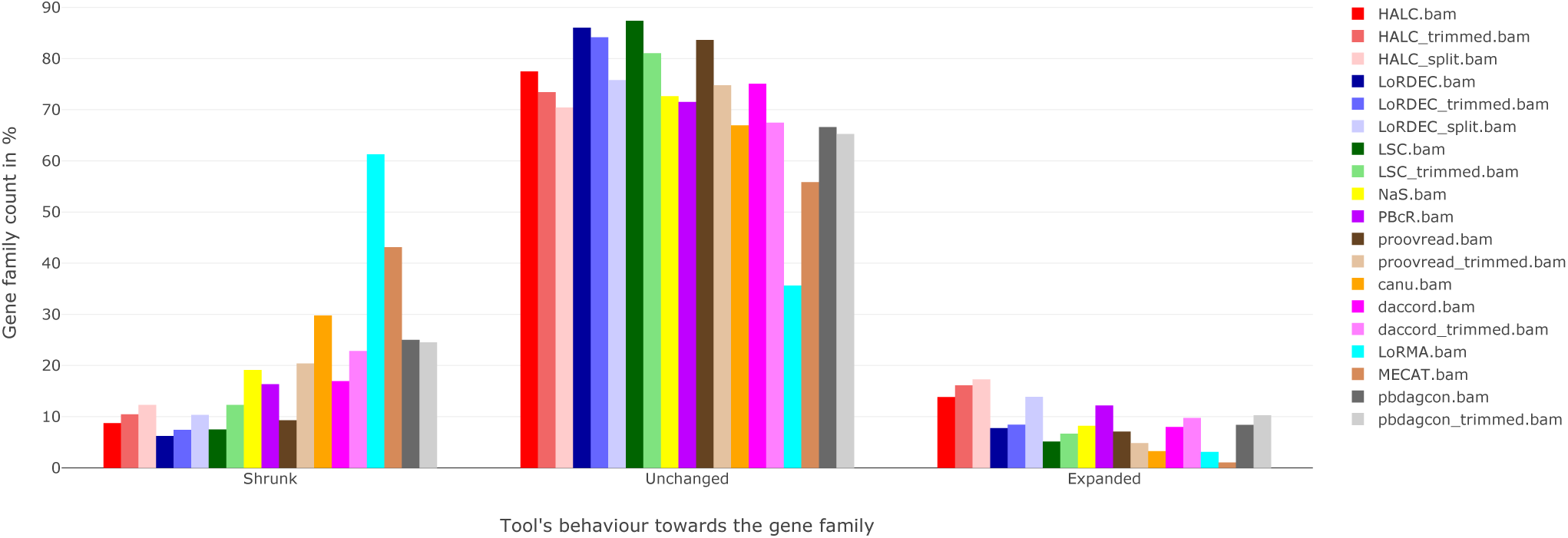
Summary of gene family size changes across error-correction tools.

### 2.8 Error-correction perturbs isoform diversity

We further investigated whether error-correction introduces a bias towards the major isoform of each gene. Note that AlignQC does not directly address this question. To answer it, we computed the following metrics: number of isoforms detected in each gene before and after correction by alignment of reads to genes, coverage of lost isoforms in genes having at least 2 expressed isoforms, and coverage of the major isoform before and after correction.

#### 2.8.1 The number of isoforms varies before and after correction

Figure 3 shows the number of genes that have the same number of isoforms after correction, or a different number of isoforms (−3, −2, −1, +1, +2, +3). In this Figure, only the genes that are expressed in both the raw and the corrected reads (for each tool) are taken into consideration. The negative (resp. positive) values indicate that isoforms were lost (resp. gained). We observe that a considerable number of genes (∼1.9k for LoRDEC(s), LSC and PBcR(s), and ∼5.4k for MECAT) lose at least one isoform in all tools, which suggests that current methods reduce isoform diversity during correction. NaS(*µ*), Proovread(t), Canu, and MECAT tend to lose isoforms the most, and HALC(s), LoRDEC(s), and PBcR(s) identify the highest number of new isoforms after correction. It is however unclear whether these lost and new isoforms are real (present in the sample) or due to mapping ambiguity, as these three latter tools split corrected reads into shorter sequences that may map better to other isoforms. We observe that the effect of trimming, on the other hand, is generally slight. Overall, the number of isoforms is mostly unchanged in LoRDEC, LoRDEC(t), and LSC.

**Fig. 3.**
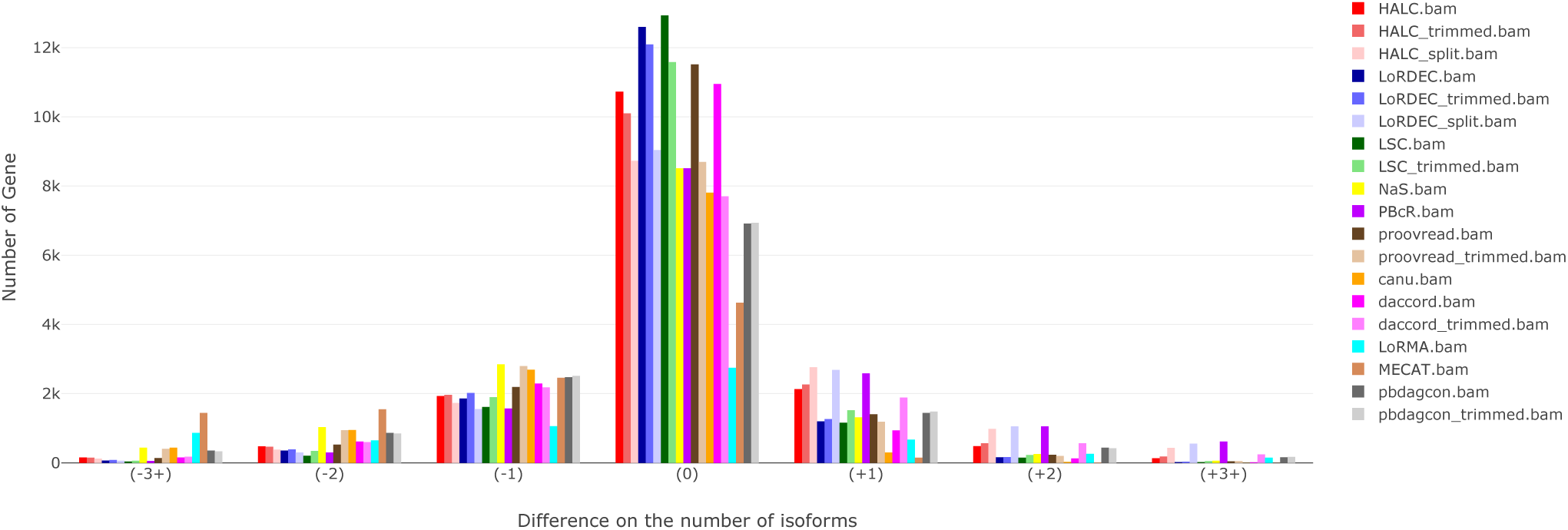
Histogram of genes having more or less isoforms after error-correction.

#### 2.8.2 Multi-isoform genes tend to lose lowly-expressed isoforms after correction

Figure 4 explores the relative coverage of isoforms that were possibly lost after correction, in genes having two or more expressed isoforms. The relative coverage of a transcript is the number of raw reads mapping to it over the number of raw reads mapping to its gene in total. Only the genes that are expressed in both the raw and the error-corrected reads (for each tool) are taken into consideration here. We anticipated that raw reads that map to a minor isoform are typically either discarded by the corrector, or modified in such a way that they now map to a different isoform, possibly the major one. The effect is indeed relatively similar across all correctors, except for MECAT, that tends to remove a higher fraction of minor isoforms, and LoRDEC and LSC, that tend to be the most conservatives. We can also note that trimming and splitting reads increase even further isoform losses in all tools, except for pbdagcon. This can be explained by the fact that lowly-expressed isoforms possibly share regions (e.g. common exons) with highly-expressed isoforms, and these shared regions are usually better corrected than regions that are unique to the lowly-expressed isoforms. If read splitting then takes places, such unique regions will then be removed from the output. Even if there are variations between a highly-expressed isoform ℐ and a lowly-expressed isoform *i*, if these variations are relatively small (e.g. a small exon skipping) and are flanked by long shared regions, it is probable that the methods will truncate the variation, correcting the unique fragment of *i* into ℐ, and potentially losing the signal that *i* is expressed (this is explored in details in Section 2.8.4). This result suggests that current error-correction tools overall do not conservatively handle reads that belong to low-expression isoforms.

**Fig. 4.**
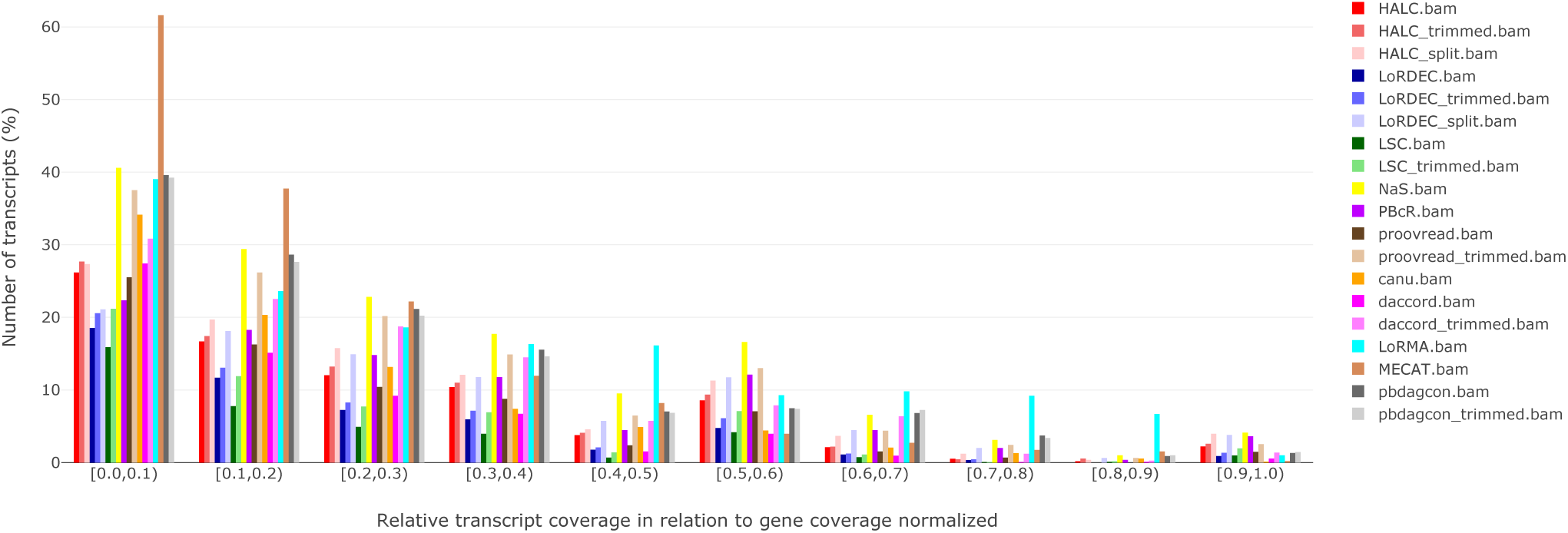
Histogram of isoforms that are lost after correction, in relation to their relative transcript coverage, in genes that have 2 or more isoforms. The y axis reflects the percentage of isoforms lost in each bin. Absolute values can be found in the Supplementary Material Figure S3.

#### 2.8.3 Minor isoforms are corrected towards major isoforms

We define a **major isoform** of a gene as the isoform with the highest coverage of that gene in the raw dataset, all other isoforms are considered to be minor. To follow-up on the previous subsection, we investigate whether correctors tend to correct minor isoforms towards major isoforms. We do so by comparing the difference of coverage of the major and the minor isoforms before and after correction. In Figure 5, we observe that the coverage of the major isoform generally slightly increases after correction. The exceptions are tools that split reads (HALC(s), LoRDEC(s), PBcR(s), and LoRMA(s)), where the coverage is increased significantly, and MECAT, where the coverage decreases significantly, likely due to a feature of MECAT’s own correction algorithm. Since these 5 tools seem to heavily distort the coverage of isoforms due to aggressive splitting or filtering steps, we will focus now on the 14 other results. The slight increase of a transcript coverage after correction is expected, as already discussed in Section 2.6: uncorrected reads that were unmapped can become mappable after correction. Therefore, the effect presented in Figure 5 could be simply due to reads being corrected to their original respective isoforms, instead of correction inducing a switch from a minor isoform to the major isoform. To verify this hypothesis, Supplementary Material Figure S4 shows that the coverage of the minor isoforms usually *decreases* after correction (*R*^2^*∈* [0.5; 0.8]), except for: i) tools that split reads (HALC(s), LoRDEC(s), PBcR(s), LoRMA(s)), which skews even more the coverage of minor isoforms, and ii) both HALC and HALC(t). This indicates that error-correction tools tend to correct reads towards the major isoforms. It is worth noting that the increase of the coverage of the major isoform is not pronounced. This is expected, as the sum of the expression of the minor isoforms is, by nature, a small fraction of the total gene expression. On the other hand, the correlation of the coverage of the minor isoforms before and after correction are far more spurious, suggesting a stronger effect. It is noteworthy that correction biases with respect to the major isoform do not appear to be specific to self-correctors nor to hybrid correctors, but an effect that happens in both types of correctors.

**Fig. 5.**
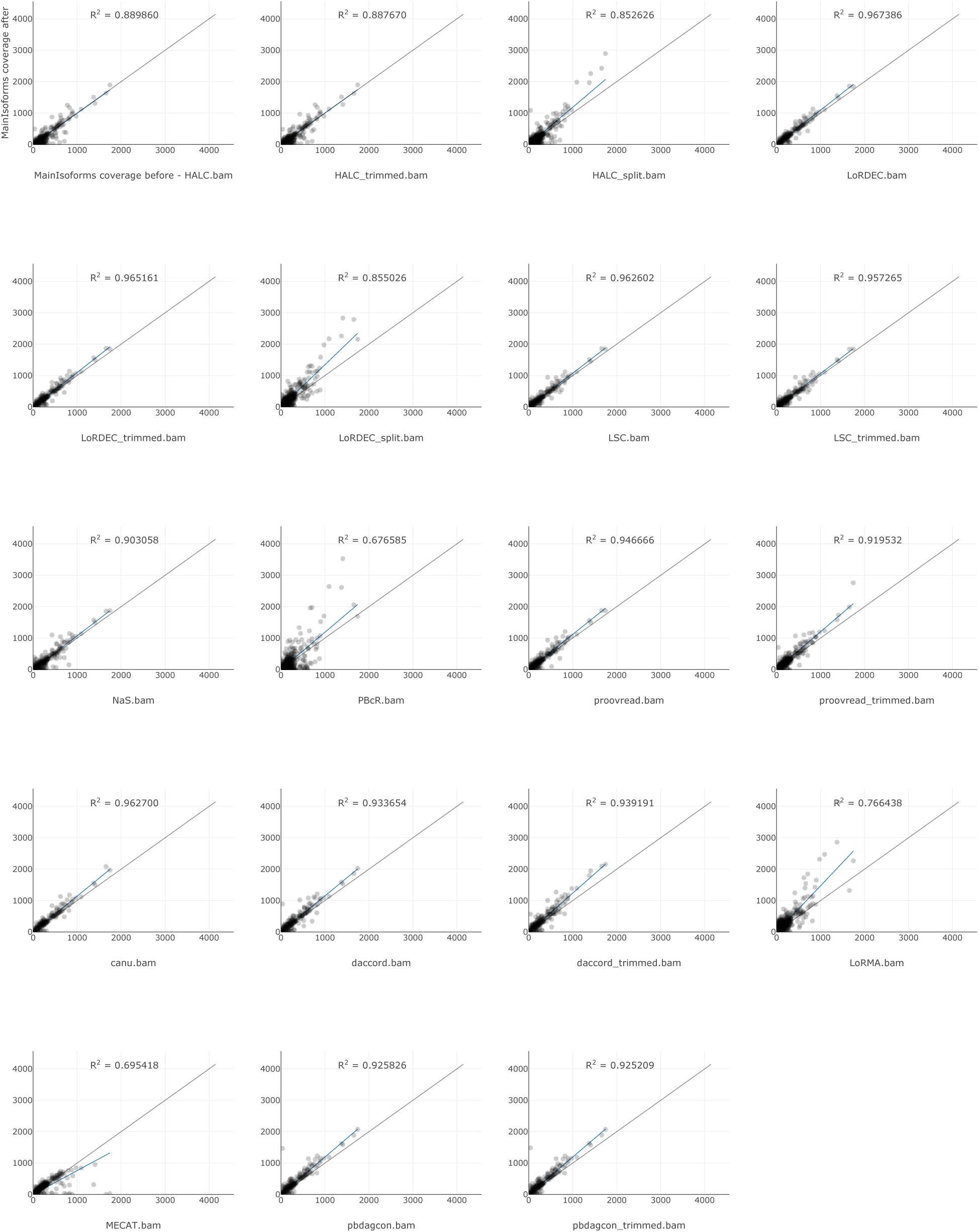
Coverage of the major isoform of each gene before and after error-correction. The x-axis reflects the number of reads mapping to the major isoform of a gene before correction, and the y-axis is after correction. Blue line: regression, black line: x=y.

#### 2.8.4 Correction towards the major isoform is more prevalent when the alternative exon is small

In order to observe if particular features of alternative splicing have an impact on error-correction methods, we designed a simulation over two controlled parameters: skipped exon length and isoform relative expression ratio. Using a single gene, we created a mixture of two simulated alternative transcripts: one constitutive, one exon-skipping. Several simulated read datasets were created with various relative abundances between major and minor isoform (in order to model a local differential in splicing isoform expression), and sizes of the skipped exon. Due to the artificial nature and small size of the datasets, many of the error-correction methods could not be run. We thus tested these scenarii on a subset of the correction methods.

In Figure 6, we distinguish results from hybrid and self-correctors, presented with respectively 100× coverage of short reads and 100x coverage of long reads, and only 100× coverage of long reads. Results on more shallow coverage (10x) and impact of simulation parameters on corrected reads sizes are presented in Supplementary Material Sections S7 and S8. Overall, hybrid correctors are less impacted by isoform collapsing than self-correctors. LoRDEC shows the best capacity to preserve isoforms in presence of alternatively skipped exons. Thus, regardless of the abundance of inclusion reads in the dataset to be corrected, 99% of reads from inclusion are corrected to inclusion form for an exon size of 10, and 100% of reads from inclusion are corrected to inclusion form for exon sizes of 50 and 100. However with less coverage, *e.g.* due to low-expressed genes and rare transcripts, all tools tend to mis-estimate the expression of isoforms (see Supplementary Material Sections S7 and S8). Self-correctors generally have a minimum coverage threshold (only daccord could be run on the 10x coverage dataset of long reads, with rather erratic results, see Supplementary Material Section S8). Even with higher coverage, not all correctors achieve to correct this simple instance. Among all correctors, only LoRDEC seems to report the expected number of each isoforms consistently in all scenarii. We could not derive any clear trend concerning the relative isoform ratios, even if the 90% ratio seems to be in favor of overcorrection towards the major isoform. Skipped exon length seems to impact both hybrid and self-correctors, small exons being a harder challenge for correctors.

**Fig. 6.**
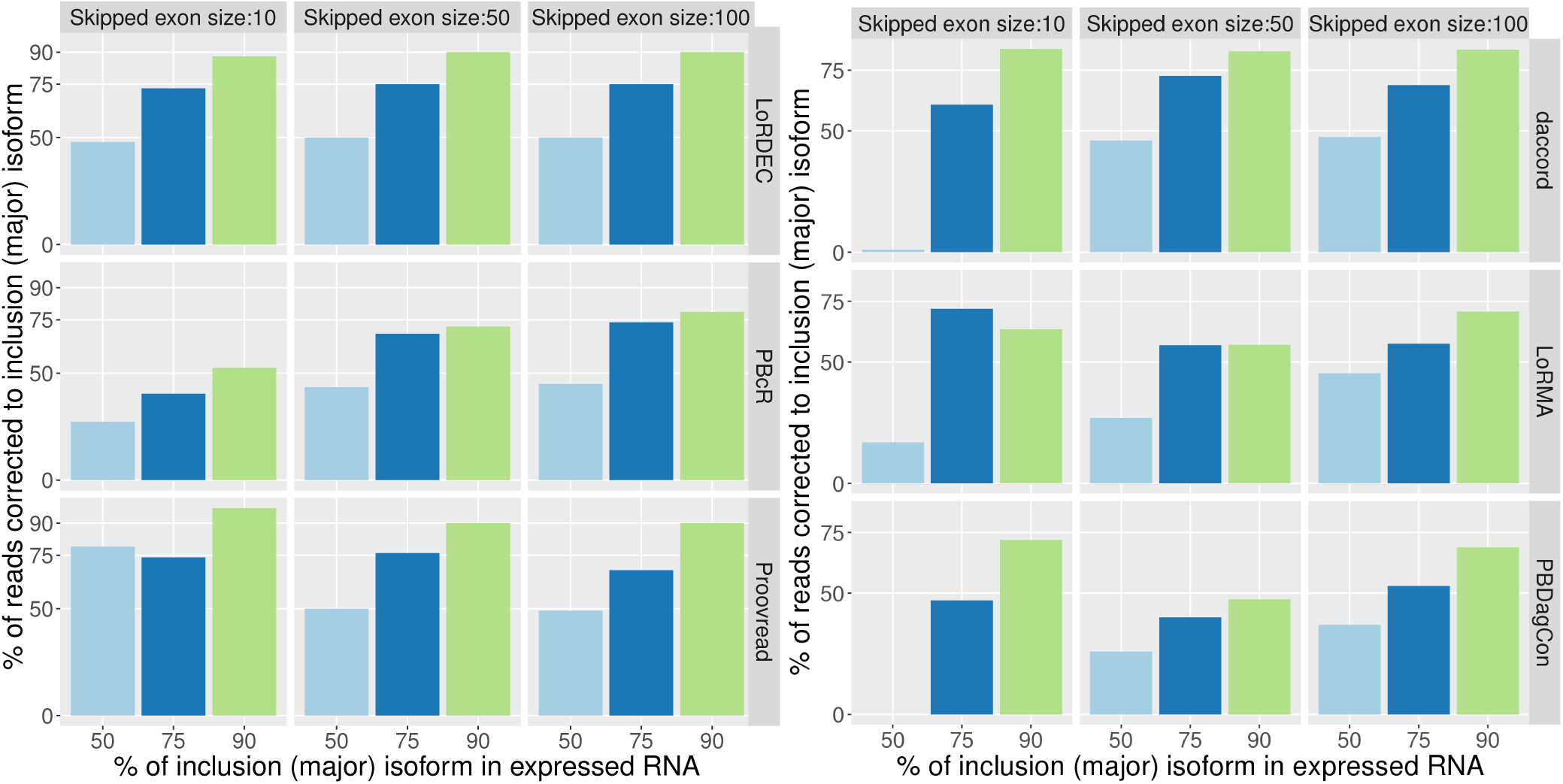
Mapping of simulated raw and error-corrected reads to two simulated isoforms, and measurements of the percentage of reads mapping to the major isoform. The two isoforms represent an alternatively skipped exon of variable size: 10 bp, 50 bp, 100bp. Left: isoform structure conservation using 100X short reads coverage and 10X long reads, using three error-correction programs, one per row: LoRDEC, PBcR, and proovread. Right: same with three self-correctors and 100X long reads: daccord, LoRMA and pbdagcon. Columns are alternative exon sizes. Bars are plots for each isoform ratio (50%; 75% and 90%) on the x-axis. On the y-axis, the closer a bar is to its corresponding ratio value on the x, the better. For instance, the bottom left light blue bar corresponds to a 50% isoform ratio with an exon of size 10, and we do not retrieve a 50% ratio after correction with Proovread (the bar does not go up to 50% on the vertical axis, but around 75% instead). The same layout applies to the right plot, where self-correctors are presented.

### 2.9 Error-correction affects splice site detection

The identification of splice sites from RNA-seq data is an important but challenging task Kaisers et al. [2017]. When mapping reads to a (possibly annotated) reference genome, mapping algorithms typically guide spliced alignments using either a custom scoring function that takes into account common splices sites patterns (e.g. GT-AG), and/or a database of known junctions. With long reads, the high error rate makes precise splice site detection even more challenging, as indels (see Section 2.5) confuse aligners, shifting predicted spliced alignments away from the true splice sites.

In this section, we evaluate how well splice sites are detected before and after error-correction. Figure 7 shows the number of correctly and incorrectly mapped splice sites for the raw and corrected reads, as computed by AlignQC. One would expect that a splice site is correctly detected when little to no errors are present in reads mapping around it. Thus, as expected, the hybrid error correction tools present a clear advantage over the non-hybrid ones, as they better decrease the per-base error rate. In the uncorrected reads, 27% of the splice sites were incorrectly mapped, which is brought down to less than 1.2% in 8 hybrid corrected outputs: HALC, HALC(t), HALC(s), LoRDEC(s), NaS(*µ*), PBcR(s), Proovread and Proovread(t). Notably, Proovread(t) presented only 0.28% incorrectly mapped splice sites. LoRDEC (2.43%) and LoRDEC(t) (2.12%) presented higher rates, but still manageable, but LSC (7.27%) and LSC(t) (5.68%) underperformed among the hybrid correctors. Among self-correction tools, LoRMA presented the lowest proportion of incorrectly detected splice sites (3.04%), however it detects ∼6.7 times less splice sites (∼280k) than the raw reads (∼1.9M), due to read splitting. The other non-hybrid tools incorrectly detected splice sites at a rate between 5.61% (daccord(t)) and 11.95% (Canu). It is worth noting that trimming usually decreased the proportion of incorrectly mapped splice sites, with a very slight impact on the total amount of identified splice sites. On the other hand, the three tools with lowest number of identified splice sites output split reads (LoRDEC(s), PBcR(s), and LoRMA(s)), identifying less than ∼1.1M splicing sites, compared to the ∼1.9M in the raw reads, and thus not being adequate for splice sites analyses. Additional detailed plots on incorrectly mapped splice sites can be found in the Supplementary Material Section S9.

**Fig. 7.**
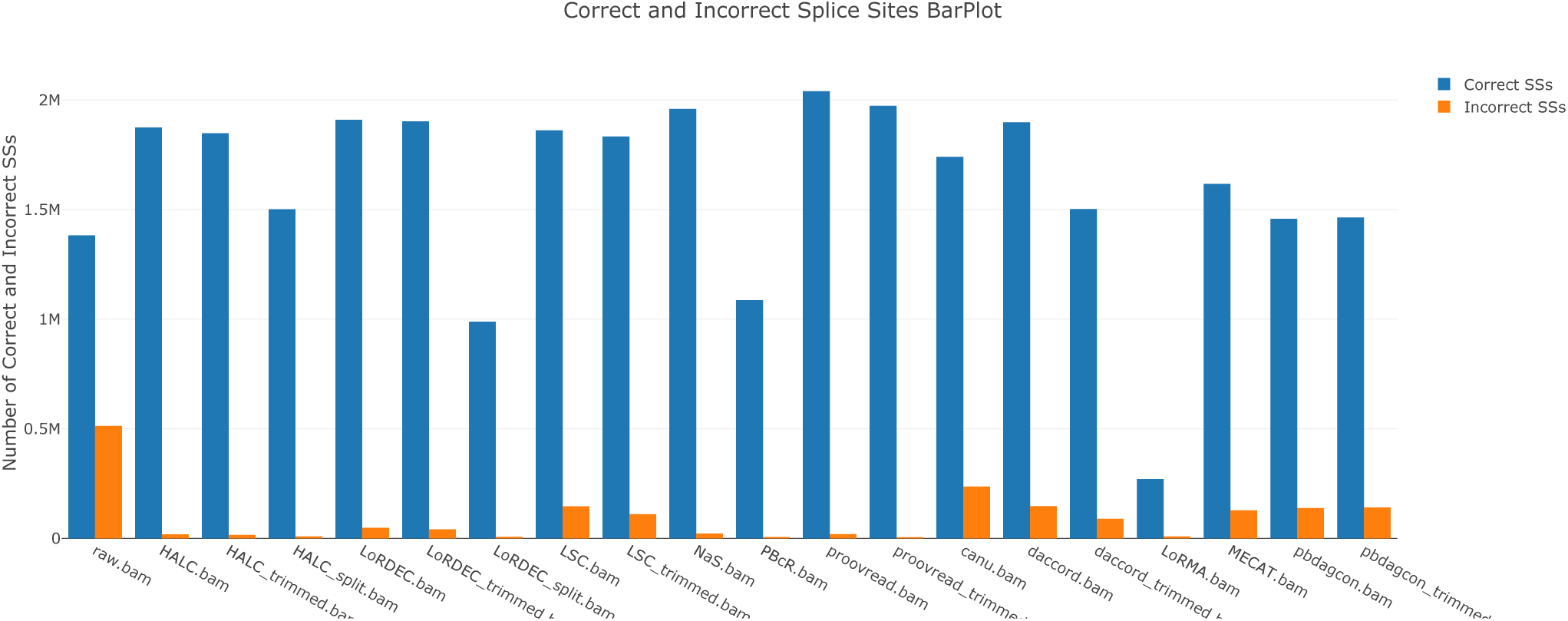
Statistics on the correctly and incorrectly mapped splice sites (abbreviated SSs) for the uncorrected (raw) and corrected reads.

### 2.10 Running time and memory usage of error-correction tools

Table 5 shows the running time and memory usage of all evaluated tools, measured using `GNU time`. The running time shown is the elapsed wall clock time (in hours) and the memory usage is the maximum resident set size (in gigabytes). All tools were ran with 32 threads. Overall, all tools were able to correct the dataset within 7 hours, except for LSC, NaS, PBcR, and Proovread, which took 63-116 hours, but also achieved some of the lowest post-correction error rates in Table 2 (except for LSC). In terms of memory usage, all tools required less than 10 GB of memory except for HALC, PBcR, Proovread and LoRMA, which required 53-166 GB. It is worth noting, however, that hybrid error correctors have to process massive Illumina datasets, which contributes to them taking higher CPU and memory usage for correction.

**Table 5.** Running time and memory usage of error-correction tools on the 1D run RNA-seq dataset

|  | HALC | LoRDEC | LSC | NaS<sup>1</sup> | PBcR<sup>2</sup> | Proovread | Canu | daccord<sup>3</sup> | daccord trimmed<sup>3</sup> | LoRMA<sup>4</sup> | MECAT | pbdagcon<sup>3</sup> | pbdagcon<sup>3</sup> trimmed |
| --- | --- | --- | --- | --- | --- | --- | --- | --- | --- | --- | --- | --- | --- |
| Running time | 3.6h | 2.4h | 99.5h | 63.2h | 116h | 107.1h | 0.7h | 6.9h<br>(7.4h) | 6.6h<br>(7.1h) | 3.4h | 0.3h | 5.7h<br>(6.2h) | 5.6h<br>(6.1h) |
| Memory usage | 60GB | 5.6GB | 2.7GB | 3GB | 166.5GB | 53.6GB | 2.2GB | 6.9GB<br>(27.2GB) | 6.8GB<br>(27.2GB) | 79GB | 9.9GB | 6.4GB<br>(27.2GB) | 6.4GB<br>(27.2GB) |

<sup>1</sup>NaS was ran in batches on a different system (TGCC cluster) than other tools; total running time was estimated based on subset of batches.
<sup>2</sup>PBcR was ran on a machine different from the others.
<sup>3</sup>daccord and pbdagcon need DAZZ.DB and DALIGNER to be ran before performing their correction. DAZZ.DB execution time and memory usage was disregarded due to being negligible. DALIGNER, however, took 0.5h and 27.2Gb of RAM. The runtime in parenthesis denotes the runtime of the tool plus DALIGNER. The memory usage in parenthesis denotes the maximum memory usage between the tool and DALIGNER.
<sup>4</sup>LoRMA was using more than its allocated 32 cores in some (short) periods of time during the run.

### 2.11 Using a different read aligner mildly but not significantly affects the evaluation

We chose GMAP (version 2017-05-08 with parameters -n 10) Wu and Watanabe [2005] to perform long reads mapping to the Ensembl r87 Mus Musculus unmasked reference genome in our analysis, since Križanović et al. [2018] show it produces the best alignment results between five alignment tools. The GMAP parameters are those from the original AlignQC publication Weirather et al. [2015]. However, Minimap2 Li [2018] is not evaluated in Križanović et al. [2018], and it is also widely used, being the default long-read mapper in several studies. In this subsection, we verify to which extent the differences between GMAP and Minimap2 can influence our evaluation. To try to highlight such differences, we chose some correctors with the worst and best performances in some measures presented in the previous analysis (made with GMAP). We thus further mapped with Minimap2 (version 2.14-r883 with parameters -`ax splice`) the following read datasets: i) raw reads; ii) LoRDEC, the corrector with the least proportion of mapped reads, but which preserves well the transcriptome diversity; iii) LoRMA(s), the tool with the highest ratio of mapped reads and number of reads, but with the lowest mean reads length, number of detected genes, and number of bases; iv) Proovread(t), the method with the lowest error rate, and highest rate of correctly mapped splice sites; v) Canu, the corrector with the highest error rate and lowest rate of correctly mapped splice sites.

We present in Table 6 the main differences between Minimap2 and GMAP on the aforementioned correctors. We can note that Minimap2 is able to map more reads than GMAP across all tools but Proovread(t). However, when both mappers are able to map almost all reads (≥98.9%), the mapping rate difference between them is low (*<*1%). The largest differences is when both mappers do not perform well, mapping less than 90% of the reads. In this case, Minimap2 does a better job, mapping 3.3% and 1.9% more reads than GMAP in the raw and LoRDEC reads, respectively, which is noticeable, but not high. A similar conclusion can be derived by looking at the mapped bases rate. The reported error rate is very similar between both mappers, with the largest difference being 0.32% in the raw reads. There is a noticeable difference, however, when we break down the errors by types. GMAP reported more mismatches, while Minimap2 reported more deletions and insertions. Nevertheless, the differences are only noticeable in the raw reads. The measure in which GMAP clearly outperformed Minimap2 is the number of detected genes, identifying between 453 (Proovread(t)) and 723 (LoRDEC) more genes, which can be considered significant. On the other hand, Minimap2 was considerably better than GMAP on mapping splice sites correctly. The difference between both mappers when they performed well, mapping more than 96.9% of the splice sites, was not significant (≤1.15%). The largest discrepancies can be seen on mapping the splice sites of the raw and Canu reads, with Minimap2 correctly aligning 18.49% and 6.64% more splice sites than GMAP.

**Table 6.** GMAP and Minimap2 results across a selection of metrics for the raw, LoRDEC, LoRMA(s), Proovread(t), and Canu reads. (G) indicates GMAP results, and (M2) Minimap2 results.

| Metric | raw(G) | raw(M2) | LoRDEC(G) | LoRDEC(M2) | LoRMA(s)(G) | LoRMA(s)(M2) | Proovread(t)(G) | Proovread(t)(M2) | Canu(G) | Canu(M2) |
| --- | --- | --- | --- | --- | --- | --- | --- | --- | --- | --- |
| mapped reads | 83.5% | 86.8% | 85.5% | 87.4% | 98.9% | 99.8% | 99.4% | 99.1% | 99.1% | 100.0%<sup>d</sup> |
| mapped bases<sup>a</sup> | 89.0% | 91.0% | 90.6% | 92.6% | 99.5% | 99.8% | 99.2% | 99.3% | 92.0% | 93.2% |
| error rate<sup>b</sup> | 14.11% | 13.79% | 4.67% | 4.65% | 0.33% | 0.31% | 2.9% | 2.73% | 6.2% | 5.97% |
| mismatches | 5.26% | 3.98% | 2.15% | 1.89% | 0.15% | 0.11% | 0.37% | 0.22% | 1.22% | 0.85% |
| deletions | 7.58% | 8.09% | 2.17% | 2.29% | 0.14% | 0.17% | 2.51% | 2.46% | 4.72% | 4.79% |
| insertions | 1.26% | 1.73% | 0.34% | 0.47% | 0.04% | 0.03% | 0.03% | 0.05% | 0.26% | 0.32% |
| nb detected genes | 16.8k | 16.2k | 16.8k | 16.0k | 14.6k | 14.2k | 6.8k | 6.3k | 12.4k | 11.7k |
| correctly mapped ss<sup>c</sup> | 72.97% | 91.46% | 97.56% | 98.69% | 99.71% | 99.58% | 96.92% | 98.07% | 88.03% | 94.67% |

<sup>a</sup> As reported by AlignQC. Percentage of bases aligned among mapped reads, taken by counting the M parts of CIGAR strings in the BAM file. Bases in unmapped reads are not counted.
<sup>b</sup> As reported by AlignQC, using a sample of 1 million bases from aligned reads segments.
<sup>c</sup> ss stands for splice sites.
<sup>d</sup> 100.0% is obtained due to rounding. A more precise mapped reads rate is 99.988%.

We can thus conclude that Minimap2 was slightly better on mapping reads and bases, and significantly better at correctly mapping splice sites. On the other hand, GMAP was considerably better at detecting genes. The error rate of the reads mapped by both tools are very similar, with GMAP reporting slightly more mismatches, and Minimap2 reporting slightly more deletions and insertions. However, these differences are mainly concentrated on the raw reads dataset, which is the less accurate.

The data shown and discussed in this section suggest that the main takeaway points from the results presented in the previous sections should not change when GMAP is replaced by Minimap2. The full report of the analysis of both mappers, containing further details, can be found in the support page of our method:https://leoisl.gitlab.io/LR_EC_analyser_support/.

### 2.12 Analysing human Nanopore direct RNA-sequencing data

We further analysed a human Nanopore *direct RNA*-sequencing dataset from the Nanopore-WGS-Consortium (see Section 2.2 for details). Since there was no corresponding Illumina sequencing for this dataset, we were able to evaluate only the non-hybrid error correction tools. Moreover, although LoRMA could be successfully executed, AlignQC could not process its output so we removed LoRMA from the evaluation. Table 7 presents some main statistics of non-hybrid error correction tools on the aforementioned dataset. In the rest of this section, we highlight the major differences and similarities between cDNA and direct RNA datasets, keeping in mind that they were performed on two different species (human and mouse, respectively).

**Table 7.** Statistics of non-hybrid error correction tools on one human Nanopore direct RNA-sequencing data from the Nanopore-WGS-Consortium.

|  | Raw | Canu | daccord | daccord(t) | MECAT | pbdagcon | pbdagcon(t) |
| --- | --- | --- | --- | --- | --- | --- | --- |
| nb reads | 894k | 533k | 744k | 792k | 224k | 772k | 771k |
| mapped reads | 76.0% | 99.2% | 97.5% | 98.2% | 99.2% | 98.6% | 98.5% |
| mean length | 739 | 917 | 849 | 755 | 1095 | 666 | 675 |
| nb of bases | 587M | 484M | 620M | 591M | 243M | 510M | 516M |
| error rate<sup>a</sup> | 14.61% | 7.61% | 6.03% | 5.47% | 6.36% | 8.1% | 8.26% |
| mismatches | 5.79% | 2.12% | 1.56% | 1.3% | 0.7% | 1.19% | 1.18% |
| deletions | 5.95% | 4.77% | 4.13% | 3.9% | 5.5% | 6.79% | 6.94% |
| insertions | 2.87% | 0.72% | 0.34% | 0.27% | 0.17% | 0.12% | 0.14% |
| homop. deletions | 2.18% | 2.01% | 1.84% | 1.81% | 2.24% | 2.18% | 2.22% |
| homop. insertions | 1.13% | 0.3% | 0.14% | 0.11% | 0.09% | 0.06% | 0.07% |
| nb detected genes | 14.1k | 8.8k | 12.3k | 10.8k | 6.3k | 10.0k | 10.1k |
| correctly mapped ss<sup>b</sup> | 76.95% | 87.97% | 92.52% | 93.94% | 92.86% | 91.16% | 91.14% |

<sup>a</sup> As reported by AlignQC, using a sample of 1 million bases from aligned reads segments.
<sup>b</sup> ss stands for splice sites.

In general, self-correctors discarded more reads on the direct RNA dataset than on the 1D cDNA dataset. Daccord(t) discarded the least number of reads (102k), while Canu and MECAT discarded a considerable amount of reads (361k and 670k, respectively), due to post-correction filtering. Due to our choice of parameters, the shortest reads in Canu and MECAT outputs were of lengths 101 and 100 bases, respectively. However, the removal of shorter reads explains only a fraction of the read losses, as the raw dataset contains only ∼59k reads smaller than 101 bases. GMAP mapped a smaller percentage of raw reads in the direct RNA dataset (76% vs 83.5% for the 1D cDNA dataset), but on the other hand, the mapping rate of error-corrected reads was generally higher. Notably, 97.5% of the daccord reads (resp. 98.2% for daccord(t)) were mapped, as opposed to 92.5% (resp. 94%) in the 1D cDNA dataset. The mean length of the corrected reads when compared to the mean length of the raw reads was also in general higher in the direct RNA dataset, which translated into tools having a number of output bases more similar to the number of bases in the raw reads in this dataset, except for MECAT.

The error rate of the raw direct RNA reads was 14.61%, 0.89% higher than in the raw 1D cDNA reads. As expected, the error rates in all tools were also higher in the direct RNA dataset, leading to worse results. The largest difference is with pbdagcon(t), where the error rate after correction of direct RNA reads is 2.55% higher than on the 1D cDNA dataset. The distribution of errors in the direct RNA dataset was more balanced, with mismatches and deletions having almost the same representation (around 5.85%), but insertions still being less represented (2.87%). The correction behaviour of the tools is similar across both datasets: insertion is the best corrected type of error, followed by mismatches, with satisfactory results, and deletions, in which the methods overall did a poor job. In particular, pbdagcon and pbdagcon(t) even *increased* the deletion error rate by 0.84% and 0.99%, respectively. The behaviour was similar on correcting homopolymer errors: homopolymer deletions were poorly corrected, with MECAT, pbdagcon and pbdagcon(t) not reducing the homopolymer deletion error rate at all, while homopolymer insertions were well corrected. The number of detected genes in the raw direct RNA dataset (14.1k) is less than in the raw 1D cDNA dataset (16.8k), although this is consistent with the difference in human/mouse genes count. Moreover, the tools also lose more genes in the direct RNA dataset. In particular, MECAT loses 6.4k genes in the 1D cDNA dataset, and 7.8k in the direct RNA dataset. The rate of correctly mapped splice sites was slightly higher in the raw direct RNA reads (76.95% vs 72.94%), but in the error corrected reads, this rate was highly similar (the largest difference was 0.44% in the daccord(t) correction).

As a result of our evaluation, and in accordance with the cDNA analysis, care should be taken when applying self-correctors to remove errors from Nanopore direct RNA-seq data. For example, Canu and MECAT present the undesirable side effect of discarding a lot of input reads, thus reducing the amount of information in the long reads, and decreasing considerably the number of detected genes. Although the tools perform well at correcting mismatches and insertions, they have trouble correcting deletions. In particular, MECAT, pbdagcon, and pbdagcon(t) perform rather poorly, with the last two even increasing the deletion error rate. Nonetheless, all tools manage to decrease the error rate so that the large majority of reads are mappable, increasing the proportion of mapping from 76% in the raw reads to at least 97.5%. If very low error-rate (¡1%) must be achieved due to requirements of a downstream application, then it seems that using hybrid correction tools is a must.

The full report of the analysis output by our method on this dataset, containing further details, can be found at https://leoisl.gitlab.io/LR_EC_analyser_support/.

## 3 DISCUSSION

This work shed light on the versatility of long-read DNA error-correction methods, which can be successfully applied to error-correction of RNA-sequencing data as well. In our tests, error rates can be reduced from 13.7% in the original reads down to as low as 0.3% in the corrected reads. This is perhaps an unsurprising realization as the error-correction of RNA-sequencing data presents similarities with DNA-sequencing, however a collection of caveats are described in the Results section. Most importantly, the number of genes detected by alignment of corrected reads to the genome was reduced significantly by several error-correction methods, mainly the non-hybrid ones. Furthermore, depending on the method, error-correction results have a more or less pronounced bias towards correction to the major isoform for each gene, jointly with a loss of the most lowly-expressed isoforms. We provided a software that enables automatic benchmarking of long-read RNA-sequencing error-correction software, in the hope that future error-correction methods will take advantage of it to avoid biases.

Detailed error-rate analysis showed that while hybrid correctors have lower error rates than self-correctors, the latter achieved comparable performance to the former in correcting substitutions and insertions. Deletions appear to be caused by systematic sequencing errors, making them fundamentally hard (or even impossible) to address in a self-correction setting. Moreover PBcR, NaS, and Proovread are one of the most resource-intensive error-correction tools, but also are some of the few correctors able to reduce the base error rate below 0.7%. The only exception to this is HALC, which presents a low running time, and ¡0.5% error rate in its split output.

We observe that hybrid correctors were able to better preserve the number of detected genes than self-correctors. The large majority of the hybrid corrections (9/12) were able to identify an amount of genes similar to the raw reads, with only NaS(*µ*), PBcR(s), and Proovread(t) being less sensitive, but still obtaining reasonable results. On the other hand, daccord was the only self-correction tool that reached the same gene identification level of the three aforementioned hybrid tools, while the others heavily truncated the transcriptome diversity. HALC, LoRDEC, LSC, Proovread (only in full-length mode) and daccord (only in full-length mode) appear to also better preserve the number of detected isoforms better than other correctors (Section 2.8.1). All tools tend to lose lowly-expressed isoforms after correction (Section 2.8.2). Several tools also tend to correct minor isoforms towards major isoforms (Section 2.8.3), mainly when the variation between them is small (Section 2.8.4). These points are expected, as most tools were mainly tailored to process DNA data where heterogeneous coverage is not expected. Furthermore, hybrid correctors outperformed self-correctors in the correction of errors near splice site junctions (Section 2.9).

As a result, we conclude that no evaluated corrector outperforms all the others across all metrics and is the most suited in all situations, and the choice should be guided by the downstream analysis, yet hybrid correction tools generally outperformed the self-correctors. For quantification, we have shown that error-correction introduces undesirable coverage biases, as per Section 2.6, therefore we would recommend avoiding this step altogether. For isoform detection, HALC, LoRDEC, LSC and Proovread (only in full-length mode) appear to be the methods of choice as they result in the highest number of detected genes in Table 2 and also preserve the number of detected isoforms as per Section 2.8. If Illumina reads are not available, then daccord (only in full-length mode) seems to be the best choice. For splice site detection, we recommend using hybrid correctors, preferably HALC, NaS, PBcR or Proovread, as per Section 2.9. The same four tools (however, HALC should be in split mode, and Proovread, in trimmed mode) are also recommended if downstream analyses require very low general error rate. Finally for all other applications, we make some general recommendations. A reasonable balance appears to be achieved by tools with no unsatisfactory values in Table 2: HALC(t), NaS(*µ*), and Proovread(t). If Illumina reads are unavailable, then the best overall self correctors seem to be daccord(t), pbdagcon and pbdagcon(t). Moreover, trimming and splitting usually increase the mapping rate and the read accuracy, but can lead to loss of information that was present in the raw reads, complicating the correct identification of genes.

Our analyses relied on a single mapping software (GMAP Wu and Watanabe [2005]) to align raw and error-corrected reads, as in previous benchmarks Weirather et al. [2017], Križanović et al. [2018]. However, we were also able to verify that Minimap2 Li [2018], another widely used mapper, produces similar results than GMAP (see Section 2.11), and thus the main messages of the analyses presented in this paper should not change by replacing GMAP by Minimap2.

As a side note, AlignQC reports that raw reads contained 1% of chimeric reads, i.e. either portions of reads that align to different loci, or to overlapping loci. The number of chimeric reads after error-correction remains in the 0.7%-1.3% range except for LoRDEC(s) (0.2%), PBcR(s) (0.1%), Proovread(t) (0.1%), LoRMA(s) (0.04%), and MECAT (0.2%), which either correctly split reads or discarded chimeric ones. We observe that HALC (4.2%), HALC(t) (3.9%), and daccord (1.7%) increased considerably the proportion of chimeric reads in the output.

Furthermore, we focused our evaluation on a single technology: Nanopore. We did an extensive analysis of 1D cDNA Nanopore data, using Illumina data for hybrid correction. We also performed a brief analysis of Nanopore direct RNA-seq data. While it would be natural to also evaluate PacBio data, we note that data from the PacBio Iso-Seq protocol is of different nature as the reads are pre-corrected by circular consensus.

In the evaluation of tools, we did not record the disk space used by each method, yet we note that it may be a critical factor for some tools (e.g. Canu) on larger datasets. We note also that genes that have low Illumina coverage are unlikely to be well corrected by hybrid correctors. Therefore our comparison does not take into account differences in coverage biases between Illumina and Nanopore data, which may benefit self-correctors. Finally, transcript and gene coverages are derived from the number of long reads aligning to a certain gene/transcript. This method enables to directly relate the results of error-correction to transcript/gene counts, but we note that in current RNA-seq analysis protocols, transcript/gene expression is still generally evaluated using short reads.

## 4 METHODS

### 4.1 Nanopore library preparation and sequencing

RNA MinION sequencing cDNA were prepared from 4 aliquots (250ng each) of mouse commercial total RNA (brain, Clontech, Cat# 636601), according to the Oxford Nanopore Technologies (Oxford Nanopore Technologies Ltd, Oxford, UK) protocol “1D cDNA by ligation (SQK-LSK108)”. The data generated by MinION software (MinKNOW 1.1.21, Metrichor 2.43.1) were stored and organized using a Hierarchical Data Format. FASTA reads were extracted from MinION HDF files using `poretools` Loman and Quinlan [2014]. We obtained 1,256,967 Nanopore 1D reads representing around 2 Gbp of data with an average size of 1650 bp and a N50 of 1885 bp. These reads were then filtered *in silico* to remove mtRNA and rRNA using BLAT Kent [2002] and `est2genome` Mott [1997], obtaining 740,776 long reads.

### 4.2 Illumina library preparation and sequencing

RNA-Seq library preparations were carried out from 500 ng total RNA using the TruSeq Stranded mRNA kit (Illumina, San Diego, CA, USA), which allows mRNA strand orientation (sequence reads occur in the same orientation as anti-sense RNA). After quantification by qPCR, each library was sequenced using 151 bp paired end reads chemistry on a HiSeq4000 Illumina sequencer.

### 4.3 Reference-based evaluation of long read error correction

A tool coined LR EC analyser, available at https://gitlab.com/leoisl/LR_EC_analyser, was developed using the Python language to analyze the output of long reads error correctors. The required arguments are the BAM files of the raw and corrected reads aligned to a reference annotated genome, as well as the reference genome in Fasta file format and the reference annotation in GTF file format. A file specifying the paralogous gene families can also be provided if plots on gene families should be created. In our main analysis, gene families were computed by selecting all paralogs from Ensembl r87 mouse genes with 80%+ identity. Note that paralogs from the same family may have significantly different lengths, and no threshold was applied with respect to coverage. The complete selection procedure is reported here: https://gitlab.com/leoisl/LR_EC_analyser/blob/master/GettingParalogs.txt. The main processing of our method involves running the AlignQC software Weirather et al. [2017] (https://github.com/jason-weirather/AlignQC) on the input BAMs and parsing its output to create custom plots. It then aggregates information into a HTML report. For example, Tables 2-4 are compilations from AlignQC results, as well as Tables 6 and 7, and Figure 7. Figures 1-5 were created processing text files built by AlignQC called “Raw data” in their output. In addition, an in-depth gene and transcript analysis can be performed using the IGV.js library (https://github.com/igvteam/igv.js) Robinson et al. [2011], Thorvaldsdottir et al. [2012]. In this paper, we did not include all plots and tables created by the tool. To visualise the full latest reports, visit https://leoisl.gitlab.io/LR_EC_analyser_support/.

### 4.4 Simulation framework for biases evaluation

In the simulation framework of Section 2.8.4, exons length and number were chosen according to resemble what is reported in eukaryotes Sakharkar et al. [2004] (8 exons, 200 nucleotides). A skipped exon, whose size can vary, was introduced in the middle of the inclusion isoform. Skipped exon can have a size of 10, 50 or 100 nt. We also allowed the ratio of minor/major isoforms (*M/m*) to vary. For a coverage of *C* and a ratio *M/m*, the number of reads coming from the major isoform is *MC* and the number of minor isoform reads is *mC*. We chose relative abundances ratios for the inclusion isoform as such: 90*/*10, 75*/*25 and 50*/*50. All reads are supposed to represent the full-length isoform. Finally for hybrid correction input, short reads of length 150 were simulated along each isoform, with 10X and 100X coverage.

During the simulation, we produced two versions of each read. The reference read is the read that represents exactly its isoform, without errors. The uncorrected read is the one in which we introduced errors. We used an error rate and profile that mimics observed R9.4 errors in ONT reads (total error rate of ∼13%, broken down as ∼5% of substitutions, ∼1% of insertions and ∼7% of deletions). After each corrector was applied to the read set, we obtained a triplet (reference, uncorrected, corrected) read that we used to assess the quality of the correction under several criteria.

We mapped the corrected reads on both exclusion and inclusion reference sequences using a fast Smith-Waterman implementation Zhao et al. [2013], from which we obtained a SAM file. It is expected that exclusion corrected reads will map on exclusion reference with no gaps, and that a deletion of the size of the skipped exon will be reported when mapping them to the inclusion. For each read, if it could be aligned to one of the two reference sequences in one block (according to the CIGAR), then we assigned it to this reference. If more blocks were needed, we assigned the read to the reference sequence with which the cumulative length of gaps is the lowest. We also reported the ratio between corrected reads size of each isoform kind and the real expected size of each reference isoform.

## KEY POINTS

- Long-read transcriptome sequencing is hindered by high error rates that affect analyses such as the identification of isoforms, exon boundaries, open reading frames, and the creation of gene catalogues.
- This review evaluates the extent to which existing long-read DNA error correction methods are capable of correcting cDNA Nanopore reads.
- Existing tools significantly lower the error rate, but they also significantly perturb gene family sizes and isoform diversity.

### ACKNOWLEDGEMENTS

The authors thank the ASTER consortium, in particular Camille Sessegolo and Vincent Lacroix, for fruitful discussions. LL acknowledges CNPq/Brazil for the support. This work was performed using the computing facilities of the CC LBBE/PRABI and the France Génomique e-infrastructure (ANR-10-INBS-09-08).

## BIOGRAPHICAL NOTE

All authors are part of the ASTER project (ANR ASTER) with the purpose of developing algorithms and software for analyzing third-generation sequencing data.

