## Supplementary Material for "Comparative assessment of long-read error-correction software applied to RNA-sequencing data"

L. Lima *et al.*

#### S1. Error-correction tools, versions, and parameters

This section describes the dependencies, versions and parameters of error-correction tools that were considered in this study, for the sake of reproducibility. All tools, except PBcR and NaS, were ran on a machine with 45 Intel Core Processor (Broadwell) @ 1999 MHz cores and 125 Gb of RAM. PBcR was ran on a cluster machine with 40 Intel(R) Xeon(R) CPU E5-2660 v3 @ 2.60GHz cores and 252 Gb of RAM, due to its memory constraints. NaS was run on the TGCC cluster, and the time measure was not available (an estimation was made, see Table 5 in the main text). All tools were ran with 32 threads.

##### S1.1. Canu

###### S1.1.1. Version

Canu v1.6 (https://github.com/marbl/canu/releases/download/v1.6/canu-1.6.Linux-amd64.tar.xz)

###### S1.1.2. Dependencies

None

###### S1.1.3. Runs and parameters

```
canu -correct -p output.canu -d $workdir genomeSize=190136026 useGrid=false
ovsMethod=sequential corOutCoverage=all minReadLength=100 minOverlapLength=100
-maxThreads=32 -maxMemory=120g corOverlapper=minimap -nanopore-raw $NANOPORE_READS
```

This correction is called **Canu.**

Parameters observations:

1. The genome size was calculated by summing up all the transcripts in ftp://ftp.ensembl.org/pub/release-87/fasta/mus_musculus/cdna/Mus_musculus.GRCm38.cdna.all.fa.gz ;
2. The parameters were inspired by a thread in Canu’s github on how to correct cDNA nanopore reads: https://github.com/marbl/canu/issues/641 ;
3. Canu was the hardest tool to run, since it was consuming several terabytes of disk, more than we had available (~7 TB). After several tries, we decided to work around this issue by changing the overlapper Canu uses in the correction step from MHAP to minimap2 (version 2.6: https://github.com/lh3/minimap2/releases/download/v2.6/minimap2-2.6_x64-linux.tar.bz2), following this thread in Canu’s github: https://github.com/marbl/canu/issues/703 . MHAP’s disk usage is a function on the genome size and repeat content, so we hypothesize that it was using a very large amount of disk space because it could be confusing highly-expressed transcripts with repeats. This might limit the usage of Canu for correcting long transcriptomic reads.

##### S1.2. d’accord

###### S1.2.1. Version

Commit 4321085e7cc426b796371e386236783d9ecd1f80 from https://github.com/gt1/daccord

**S1.2.2. Dependencies**

1. DAZZ_DB: commit 8ab7daf27123a3dfd17189af168ebe22b343170f from https://github.com/thegenemyers/DAZZ_DB.git;
2. DALIGNER: commit 381fa920935a6bb469353cfc5c83d301579b5e04 from https://github.com/thegenemyers/DALIGNER.git;
3. Since d'accord needs PacBio headers to work, artificial PacBio headers were added to the Nanopore reads by using a script from Nanocorrect (commit 9d8d28d655783555348393e6143ddf7685d03f6a from https://github.com/jts/nanocorrect.git). The script described is https://github.com/jts/nanocorrect/blob/master/nanocorrect-preprocess.pl ;
4. libmaus2 version 2.0.420-release-20171116172420: https://github.com/gt1/libmaus2/archive/2.0.420-release-20171116172420.zip

**S1.2.3. Runs and parameters**

1. PacBio headers were added to the Nanopore reads using the Nanocorrect script (using default parameters);
2. The Dazzler Data Base was built using DAZZ_DB (module fasta2DB, using default parameters);
3. The Dazzler Data Base was split using DAZZ_DB (module DBsplit, with parameter -a);
4. DALIGNER jobs were created and run using DALIGNER (module HPC.daligner, with parameters -T32 -l200). In total, 8 .las files were created by DALIGNER.
5. d'accord was ran for each .las file produced by DALIGNER (8 in total):
   **daccord -t32 <las_file_x> <DAZZ_DB> > output.daligner.x.fasta**
   which generated 8 corrected reads file, which were concatenated. This correction is called **daccord(t)**.
6. d'accord was also ran to produce full alignments, with the -f1 parameter:
   **daccord -t32 -f1 <las_file_x> <DAZZ_DB> > output.daligner.x.fasta**
   which also generated 8 corrected reads file, which were concatenated. This correction is called **daccord**.

**S1.3. HALC**

**S1.3.1. Version**

HALC v1.1 (https://github.com/lanl001/halc/archive/v1.1.tar.gz)

**S1.3.2. Dependencies**

1. Blasr v5.3.4323a52 (https://github.com/PacificBiosciences/blasr)
2. Trinity-v2.5.1 (https://github.com/trinityrnaseq/trinityrnaseq/releases/tag/Trinity-v2.5.1)

**S1.3.3. Runs and parameters**

HALC requires contigs assembled from the corresponding short reads in FASTA format. For transcriptomic data, in its paper (https://bmcbioinformatics.biomedcentral.com/articles/10.1186/s12859-017-1610-3), Trinity got the best results, so we also used Trinity as:

```
Trinity --seqType fq --max_memory 100G --left $ILLUMINA_READS_1 --right
$ILLUMINA_READS_2 --CPU 32 --output $OUTPUT_DIR
```

HALC was run as:

```
python runHALC.py -t 32 -o $ILLUMINA_READS $NANOPORE_READS $TRINITY_CONTIGS
```

HALC outputs three corrections: full-length error-corrected reads (`*corrected.fa`), trimmed error-corrected reads (`*trim.fa`) and split error-corrected reads (`*split.fa`). These corrections are called, respectively, **HALC**, **HALC(t)**, and **HALC(s)**.

**S1.4. LoRDEC**

**S1.4.1. Version**

LoRDEC v0.6 with GATB Core v1.0.6.

**S1.4.2. Dependencies**
None

**S1.4.3. Runs and parameters**
```
lordec-correct -T 32 -i $NANOPORE_READS -2 ${ILLUMINA_READS_1},${ILLUMINA_READS_2} -o output.lordec.fasta -k 21 -s 2
```

This correction is called **LoRDEC**.
Parameters observations:
Parameters were chosen from http://www.atgc-montpellier.fr/lordec/: "For bacterial species or eukaryotic species with small genomes, you may choose k=19 or 17, and s=2 or 3. For species with larger genomes, k=21 and s=2 or 3."

For the trimmed output, we ran:
```
lordec-trim -i ${LORDEC_CORRECTION} -o ${LORDEC_TRIMMED_CORRECTION}
```
This correction is called **LoRDEC(t).**

For the split output, we ran:
```
lordec-trim-split -i ${LORDEC_CORRECTION} -o ${LORDEC_SPLIT_CORRECTION}
```
This correction is called **LoRDEC(s).**

**S1.5. LoRMA**
**S1.5.1. Version**
LoRMA v0.4 (https://www.cs.helsinki.fi/u/lmsalmel/LoRMA/LoRMA-0.4.tar.gz).

**S1.5.2. Dependencies**
None

**S1.5.3. Runs and parameters**
```
lorma.sh -threads 32 $NANOPORE_READS
```
This correction is called **LoRMA(s)**.

**S1.6. LSC**
**S1.6.1. Version**
LSC-2.0 (https://www.healthcare.uiowa.edu/labs/au/LSC/files/LSC-2.0.tar.gz)

**S1.6.2 Dependencies**
1. Bowtie2 v2.3.4.3 (https://downloads.sourceforge.net/project/bowtie-bio/bowtie2/2.3.4.3/bowtie2-2.3.4.3-linux-x86_64.zip)
2. samtools v1.6 (https://sourceforge.net/projects/samtools/files/samtools/1.6/samtools-1.6.tar.bz2/download)

**S1.6.3. Runs and parameters**
LSC was ran in 3 separated modes to allow for parallel execution (see Section "Step 3 (alternative) - Parallelize execution of the LSC command" in https://www.healthcare.uiowa.edu/labs/au/LSC/LSC_tutorial.asp). The command lines for each mode are described below.

```
#setup some parameters used in all modes
SUFFIX_COMMANDS="--samtools_path $SAMTOOLS_PATH --short_read_file_type fq
--specific_tempdir $LSC_TMPDIR --output $RUN_DIR"
```

```
#run LSC in mode 1
$TOOL_PATH --mode 1 --long_reads $NANOPORE_READS --short_reads $ILLUMINA_READS_1
$ILLUMINA_READS_2 --threads $THREADS $SUFFIX_COMMANDS

#run the tool in mode 2
BATCH_COUNT=`cat $LSC_TMPDIR/batch_count`
#create the jobs to run in parallel
echo "" > $LSC_TMPDIR/parallel_jobs
for i in `seq 1 $BATCH_COUNT`;
do
     echo "$TOOL_PATH --parallelized_mode_2 $i --threads 1 $SUFFIX_COMMANDS" >>
$LSC_TMPDIR/parallel_jobs
done
parallel -j$THREADS :::: $LSC_TMPDIR/parallel_jobs

#run the tool in mode 3
$TOOL_PATH --mode 3 $SUFFIX_COMMANDS
```

From the LSC output, we call the file `full_LR.fa` as the correction **LSC** and `corrected_LR.fa` as the correction **LSC(t)**.

**S1.7. MECAT**
**S1.7.1. Version**
Commit 3898797d5d0ead78b14af65089f6be32263ca103 from https://github.com/xiaochuanle/MECAT

**S1.7.2. Dependencies**
None

**S1.7.3. Runs and parameters**
```
mecat2pw -j 0 -d $NANOPORE_READS -o overlaps.txt -w wrk_dir -t 32 -x 1 -a 100
mecat2cns -i 0 -x 1 -t 32 -a 100 -l 100 overlaps.txt $NANOPORE_READS
output.mecat.fasta
```

This correction is called **MECAT**.
Parameters observations:
Parameters were inspired by https://github.com/xiaochuanle/MECAT#assemblying-nanopore-data

**S1.8. NaS**
**S1.8.1. Version**
Commit c8444284d5914283270615c3db967ce75cf7f9e1 from https://github.com/institut-de-genomique/NaS

**S1.8.2. Dependencies**
BLAT, Newbler, LAST, GNU parallel, and many perl modules as listed here:
https://github.com/institut-de-genomique/NaS

**S1.8.3. Runs and parameters**
Parameters:
**--mode fast --nb_proc 10  --selectReads_tool minimap --rmtmp no --ilmn_size 150  --t 2 --k 32 --untgl_seq_size 300**

This correction is called **NaS(μ)**.

##### S1.9. PBcR
###### S1.9.1. Version
Linux pre-compiled version wgs-8.3rc2-Linux_amd64 (http://sourceforge.net/projects/wgs-assembler/files/wgs-assembler/wgs-8.3/wgs-8.3rc2-Linux_amd64.tar.bz2)

###### S1.9.2. Dependencies
None

###### S1.9.3. Runs and parameters
```
PBcR -length 200 -partitions 200 -threads 32 -genomeSize 190136026 -libraryname output.PBcR.fasta -s <oxford.spec> -fastq $NANOPORE_READS $ILLUMINA.frg
```

This correction is called **PBcR(s)**.
Parameters observations:

1. The genome size was calculated by summing up all the transcripts in ftp://ftp.ensembl.org/pub/release-87/fasta/mus_musculus/cdna/Mus_musculus.GRCm38.cdna.all.fa.gz
2. The .spec file was one suited for ONT data, available in http://wgs-assembler.sourceforge.net/wiki/index.php/PBcR#Assembling_a_MinION_dataset
3. $ILLUMINA.frg was obtained by running `PBcR/wgs-8.3rc2/Linux-amd64/bin/fastqToCA -libraryname illumina -insertsize 182 64 -technology illumina -type sanger -innie -mates ${ILLUMINA_READS_1},${ILLUMINA_READS_2}`

##### S1.10. pbdagcon
###### S1.10.1. Version
Commit 8f1a59d796093fccd986375ef642b6628ff2a7db from https://github.com/PacificBiosciences/pbdagcon

###### S1.10.2. Dependencies
None

###### S1.10.3. Runs and parameters
1. The Dazzler Data Base and the .las files created to run d'accord were used as input to module dazcon of pbdagcon. dazcon was ran for each las file produced by DALIGNER (8 in total). The parameters used for dazcon were inspired by https://www.biorxiv.org/content/biorxiv/early/2017/02/06/106252.full.pdf :
   ```
   dazcon -c 2 -l 100 -m 10000 -j 32 -s <DAZZ_DB> -a <las_file_x> output.pbdagcon.x.fasta
   ```
   which generated 8 corrected reads file, which were concatenated. This correction is called **pbdagcon**.
2. pbdagcon was also ran in trimmed mode by adding parameter -t 1 to the previous command line:
   ```
   dazcon -c 2 -l 100 -m 10000 -t 1 -j 32 -s <DAZZ_DB> -a <las_file_x> output.pbdagcon.x.fasta
   ```
   The trimmed correction is called **pbdagcon(t)**.

##### S1.11. proovread
###### S1.11.1. Version
Commit 21fecc052f65a22319b36bae8d784303be691d14 from https://github.com/BioInf-Wuerzburg/proovread

**S1.11.2. Dependencies**
None

**S1.11.3. Runs and parameters**
```
proovread --long-reads=$NANOPORE_READS --short-reads=$ILLUMINA_READS_1
--short-reads=$ILLUMINA_READS_2 -t 32
```

Proovread creates two output files: proovread.trimmed.fq and proovread.untrimmed.fq, which are called **proovread(t)** and **proovread**, respectively.

**S1.12. Justifications for tools that we considered but did not run**
**S1.12.1 Nanopolish and nanocorrect**
Jared Simpson (one of the authors of nanopolish and nanocorrect) writes in http://simpsonlab.github.io/2016/02/25/deprecating-nanocorrect/ that a pipeline to process ONT data was developed, consisting of nanocorrect + Celera Assembler, and then nanopolish to compute the final consensus sequence. However, nanocorrect is said to be deprecated as the Canu and the Miniasm assemblers are clearly better at building contigs than their pipeline, finally suggesting to build contigs with Canu and using nanopolish to compute the final consensus sequence. Thus, we did not add nanocorrect to the set of error-correction tools as it is deprecated. Moreover, we decided also to not add nanopolish since it is tailored to polish a consensus, not error correct reads. As such, the input to nanopolish should be a reference-quality sequence, e.g possibly reads that were already error-corrected or that already contain a very low error rate, which is not the case of raw 1D Nanopore or CLR PacBio reads nowadays.

**S1.12.2 falcon_sense**
Canu internally uses an adapted version of falcon_sense, so we decided that the results might be too similar to be informative, and prioritized other correctors. Running other correctors could provide more interesting results and a more comprehensive study.

**S1.12.3 PBcR non-hybrid version**
PBcR was run only in hybrid mode, as the authors suggest using Canu over the non-hybrid mode.

**S1.12.4 LSCPlus**
LSCPlus (https://www.ncbi.nlm.nih.gov/pmc/articles/PMC5103424/) seemed a promising tool to run, since it is a tool designed to correct PacBio RNA-seq long reads using Illumina short reads. It is an improvement over the LSC tool, which uses the latter as a reference, but overcomes the disadvantage of LSC's time consumption and improves the correction quality. However, the software webpage (http://www.herbbol.org:8001/lscplus/) was unreachable for months, from the conception of this study until its submission.

### S2. Ratio of homopolymer errors over all errors

#### S2.1. Ratio of homopolymer deletions over all deletions

(A) Hybrid tools

| | Raw | HALC | HALC (t) | HALC (s) | LoRD EC | LoRD EC(t) | LoRD EC(s) | LSC | LSC(t) | NaS(μ) | PBcR(s) | Proov read | Proov read (t) |
|---|---|---|---|---|---|---|---|---|---|---|---|---|---|
| Ratio (%) | 39.9 | 32.9 | 29.7 | 17.6 | 35.8 | 36.4 | 48.7 | 23.5 | 21.6 | 22.2 | 33.3 | 30.5 | 22.2 |

(B) Non-hybrid tools

| | Raw | Canu | daccord | daccord(t) | LoRMA(s) | MECAT | pbdagcon | pbdagcon (t) |
|---|---|---|---|---|---|---|---|---|
| Ratio (%) | 39.9 | 51.0 | 56.0 | 62.7 | 72.5 | 50.1 | 44.7 | 43.9 |

Table S1. Ratio of homopolymer deletions over all deletions

#### S2.2. Ratio of homopolymer insertions over all insertions

(A) Hybrid tools

| | Raw | HALC | HALC (t) | HALC (s) | LoRD EC | LoRD EC(t) | LoRD EC(s) | LSC | LSC(t) | NaS(μ) | PBcR(s) | Proov read | Proov read (t) |
|---|---|---|---|---|---|---|---|---|---|---|---|---|---|
| Ratio (%) | 31.7 | 23.8 | 21.4 | 20.0 | 28.1 | 29.2 | 28.6 | 23.4 | 22.5 | 12.5 | 10.5 | 27.3 | 33.3 |

(B) Non-hybrid tools

| | Raw | Canu | daccord | daccord(t) | LoRMA(s) | MECAT | pbdagcon | pbdagcon (t) |
|---|---|---|---|---|---|---|---|---|
| Ratio (%) | 31.7 | 28.6 | 21.4 | 15.8 | 25.0 | 24.1 | 22.2 | 20.0 |

Table S2. Ratio of homopolymer insertions over all insertions

#### S3. Error-correction perturbs the number of reads mapping to the transcripts

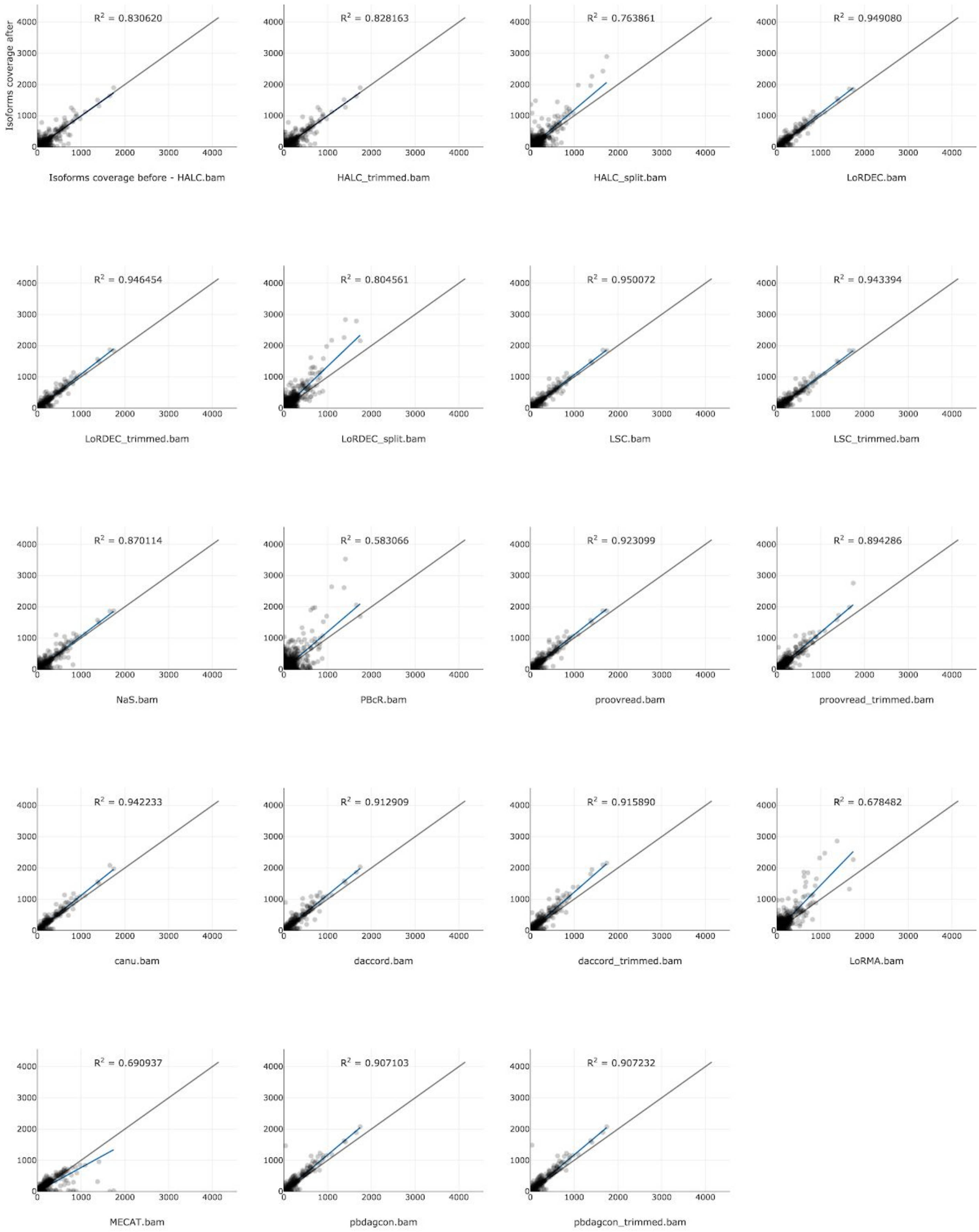


Figure S1. Number of reads mapping to isoforms ($C_T$) before and after correction for each tool. The isoforms taken into account here were expressed in either the raw dataset or after the correction by the given tool.

#### S4. Detailed view of paralogous gene families change per tool

This scatterplot shows the sizes of the paralogous gene families before and after correction, detailing what was shown in Fig. 2 of the main text. The color code of a data point is the number of gene families on that data point.

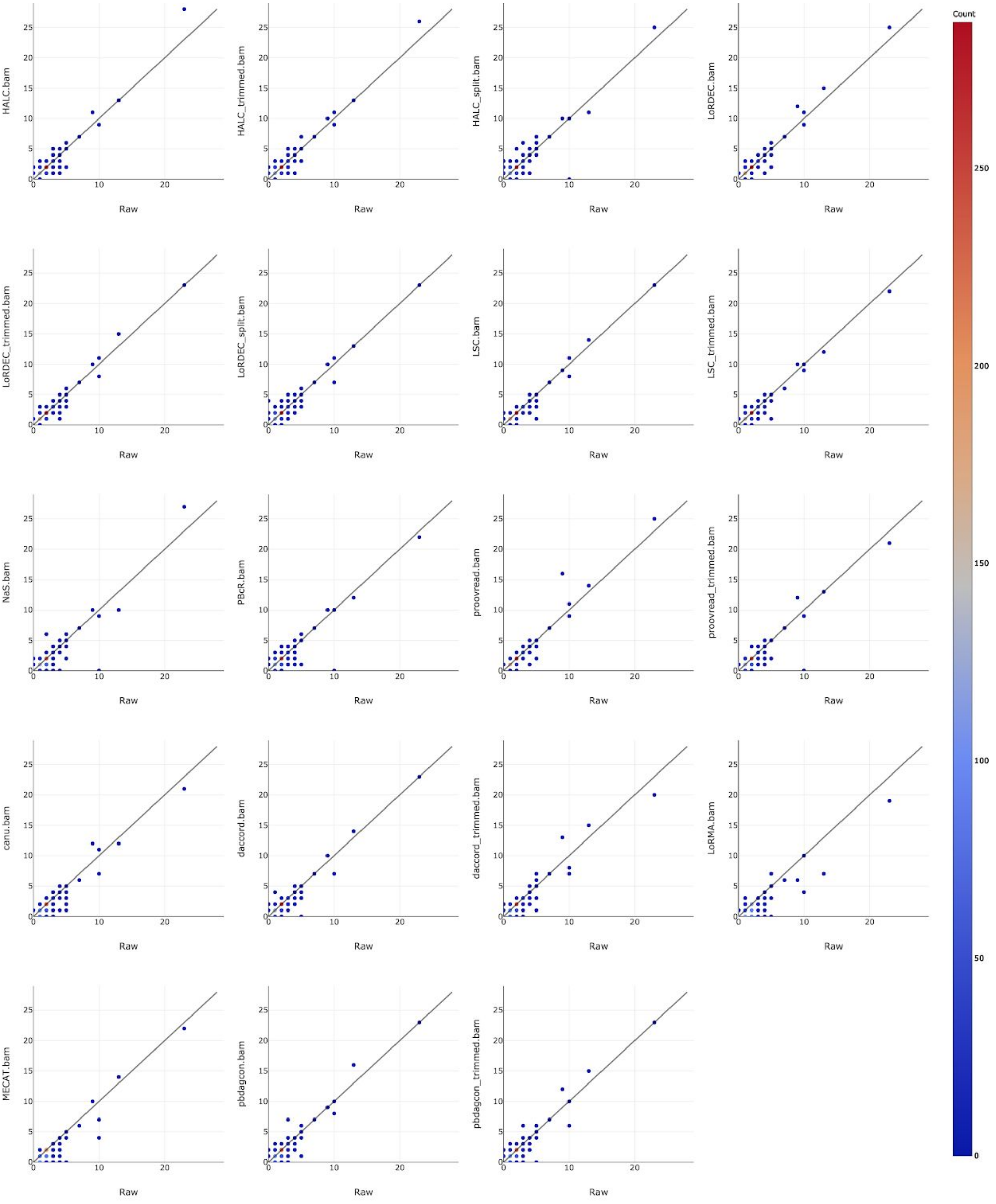

Figure S2. Sizes of paralogous gene families before and after error-correction, considering only those with cardinality at least 2. The color code of a data point is the number of gene families on that data point.

#### S5. Multi-isoform genes tend to lose lowly-expressed isoforms after correction

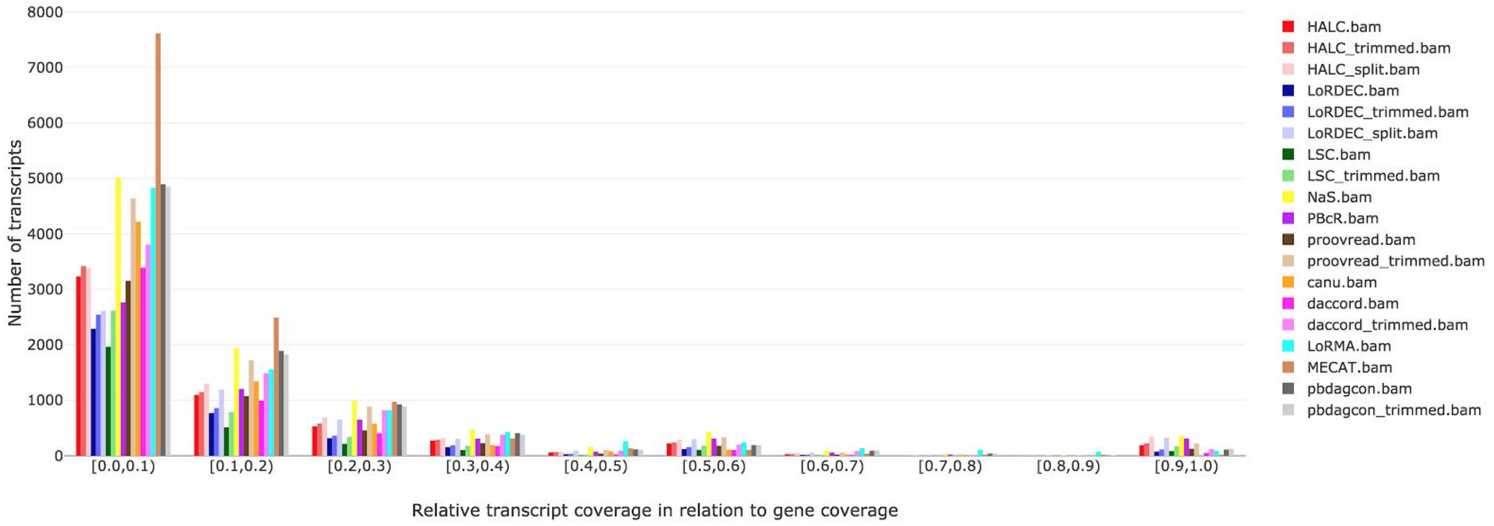


Figure S3. Absolute values corresponding to Fig 4 in the main text. Histogram of isoforms that are lost after correction, in relation to their relative transcript coverage, in genes that have 2 or more isoforms. The y axis reflects the number of isoforms.

#### S6. Minor isoforms are corrected toward major isoforms

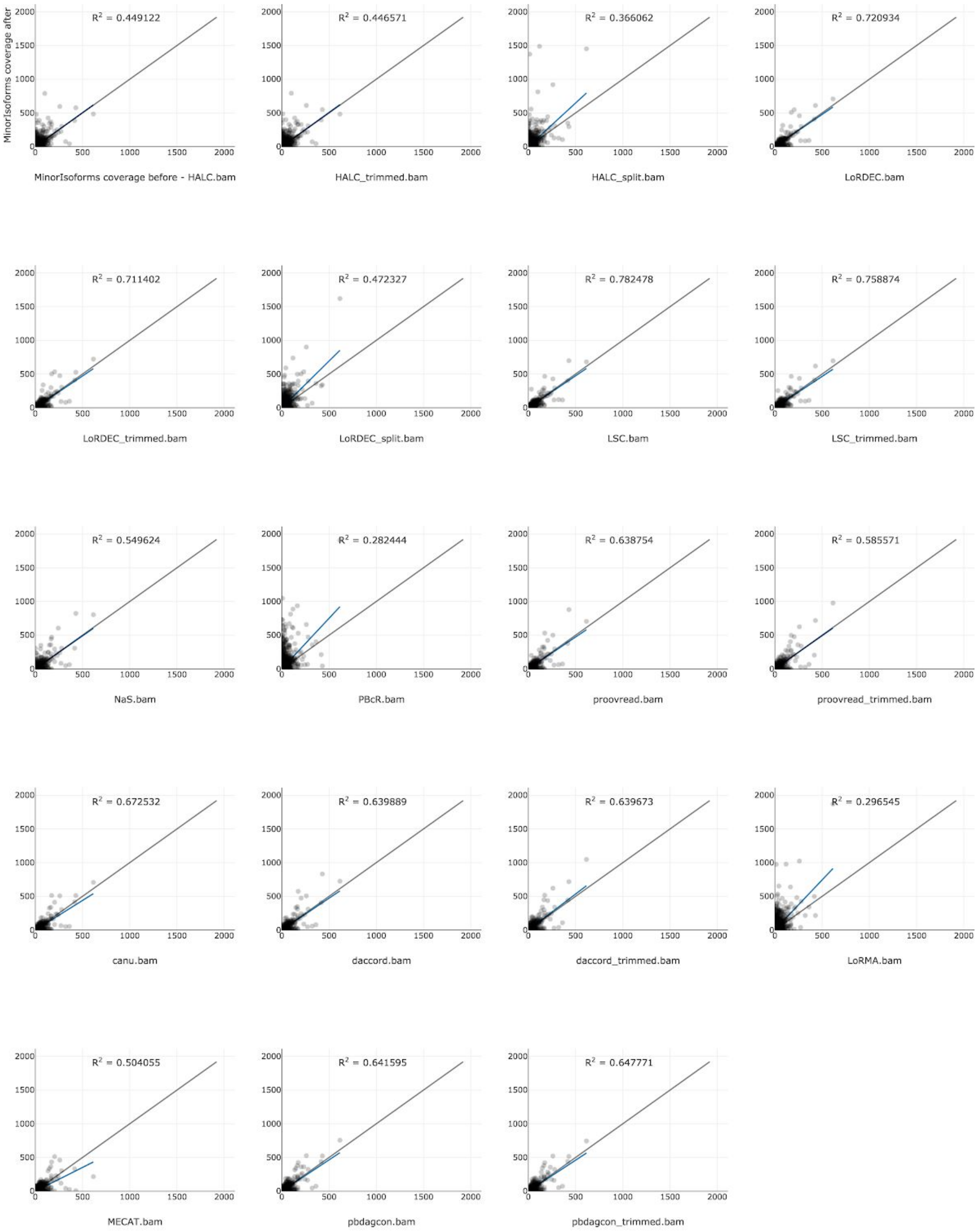


Figure S4. Coverage of the minor isoforms of each gene before and after error-correction. The x-axis reflects the number of reads mapping to the minor isoform of a gene before correction, and the y-axis is after correction. Blue line: regression, black line: x=y.

### S7. Additional results to correction towards the major isoform for hybrid correctors

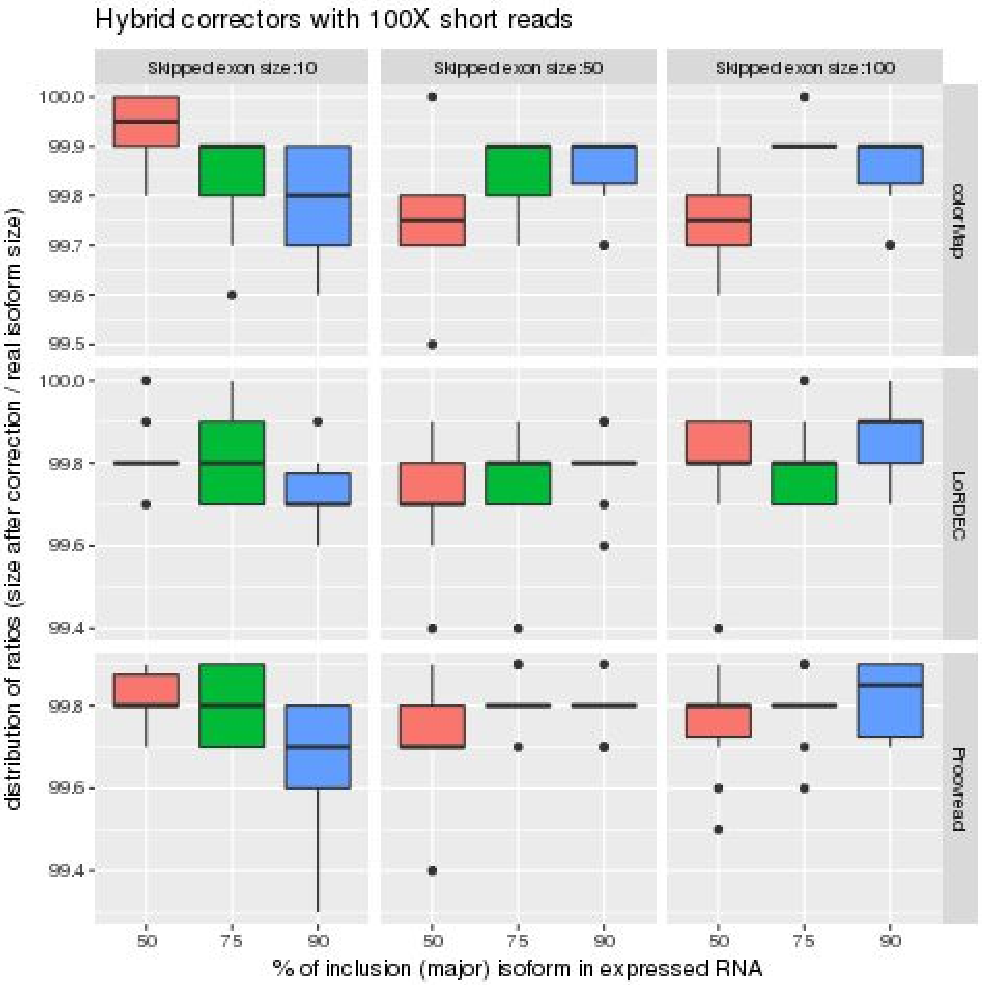


Figure S5. Distribution of ratios (size after correction/real isoform size) measured using all corrected reads, for 100X short reads. Observe in this Figure that all hybrid correctors manage to yield reads of size > 99% of the expected size.

**Impact of short read coverage on hybrid correction**

We used a 10X short reads coverage to allow a comparison with the previous results and see how correctors would behave on isoforms poorly covered.

Globally hybrid correctors make more errors when correcting to an isoform or the other. They also yield reads shorter than with 100X coverage.

LoRDEC seems to be the less sensitive to the coverage change.

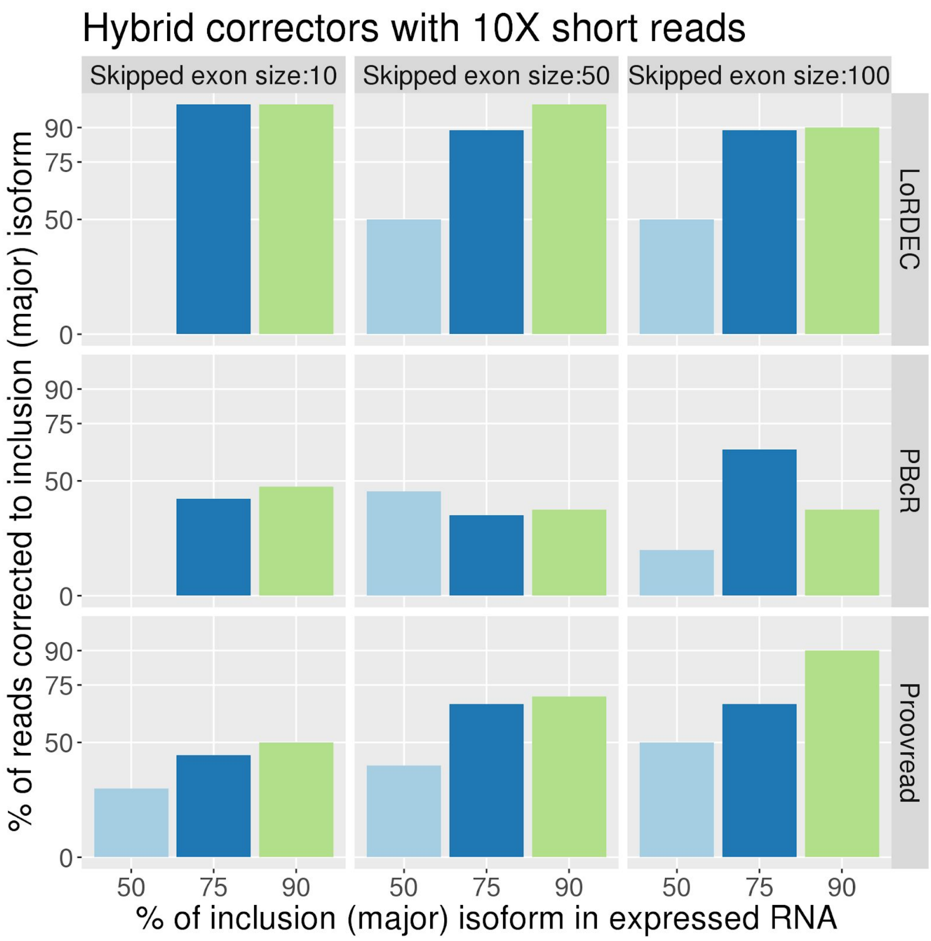


Figure S6. Counterpart of Figure 6 (from the main text) but with 10x coverage of short reads instead of 100x.

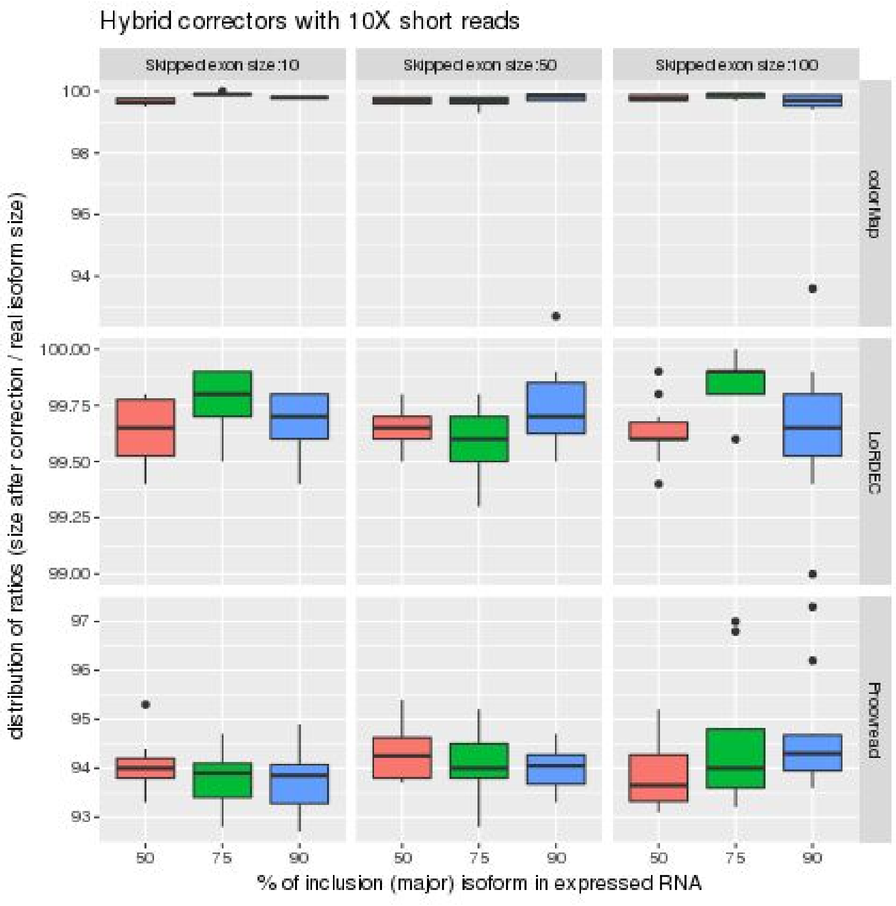


Figure S7. Counterpart of Figure S5 but with 10x coverage of short reads instead of 100x.

#### S8. Additional results to correction towards the major isoform for self correctors

Simulation was realized with 10X long reads. Again we plot % of reads corrected to inclusion (major) isoform. Shallow coverage does not allow Daccord to retrieve each isoform in its original ratio. For instance for skipped exon size of 10, all reads were corrected to exclusion isoform.
Then we plot the distribution of ratios (size after correction/ real isoform size) measured using all corrected reads, and observe that reads corrected by daccord can be way shorter than expected.

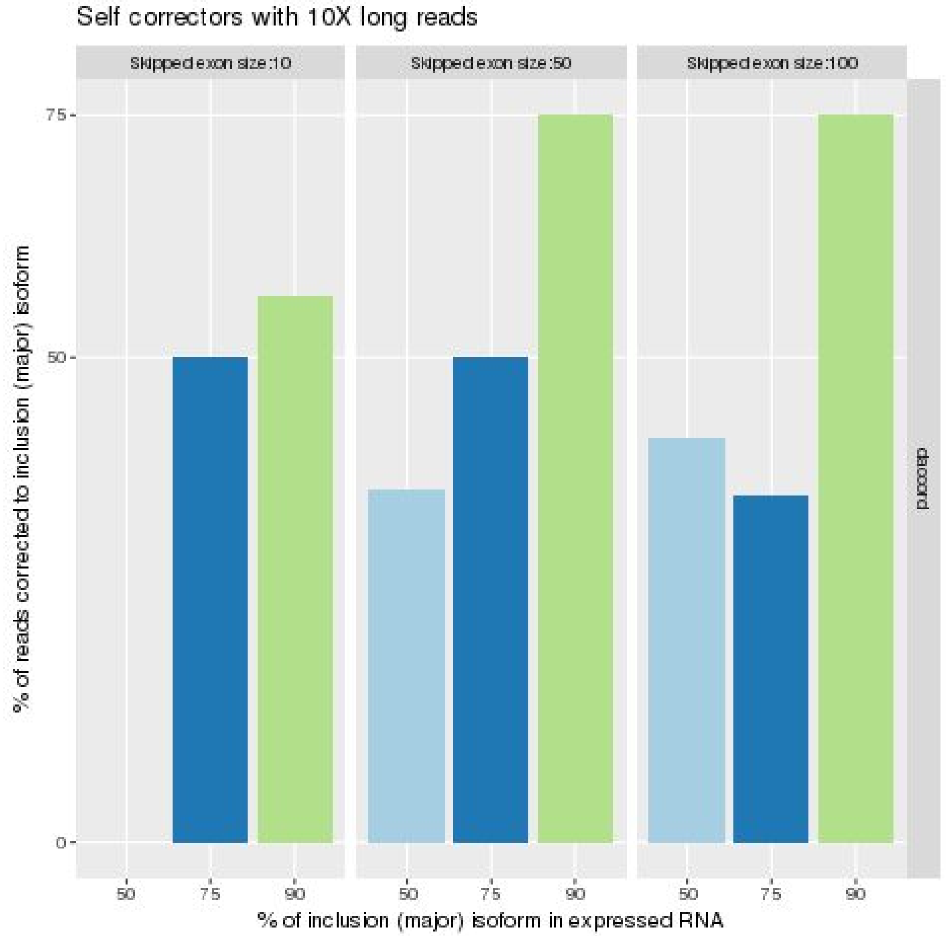


Figure S8. Counterpart of Figure 6 of the main text but with 10x coverage of long reads instead of 100x. Only daccord could be run with such a low coverage.

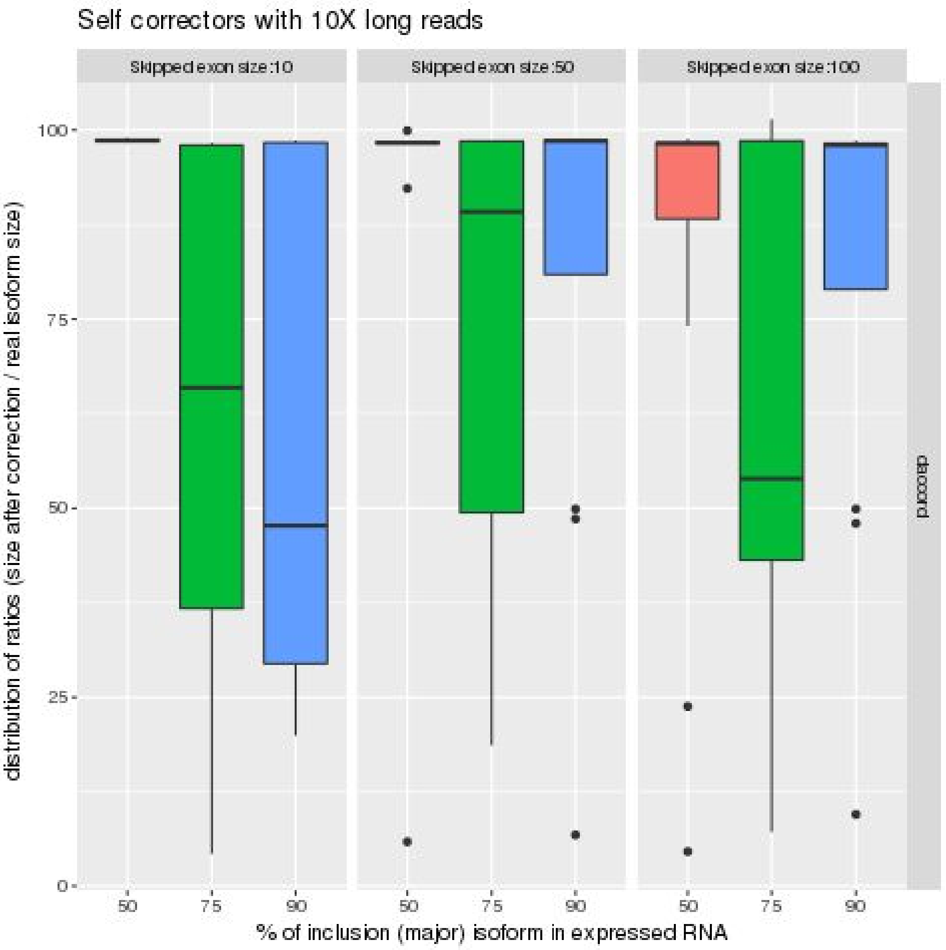


Figure S9. Counterpart of Figure S5 but with 10x coverage of long reads.

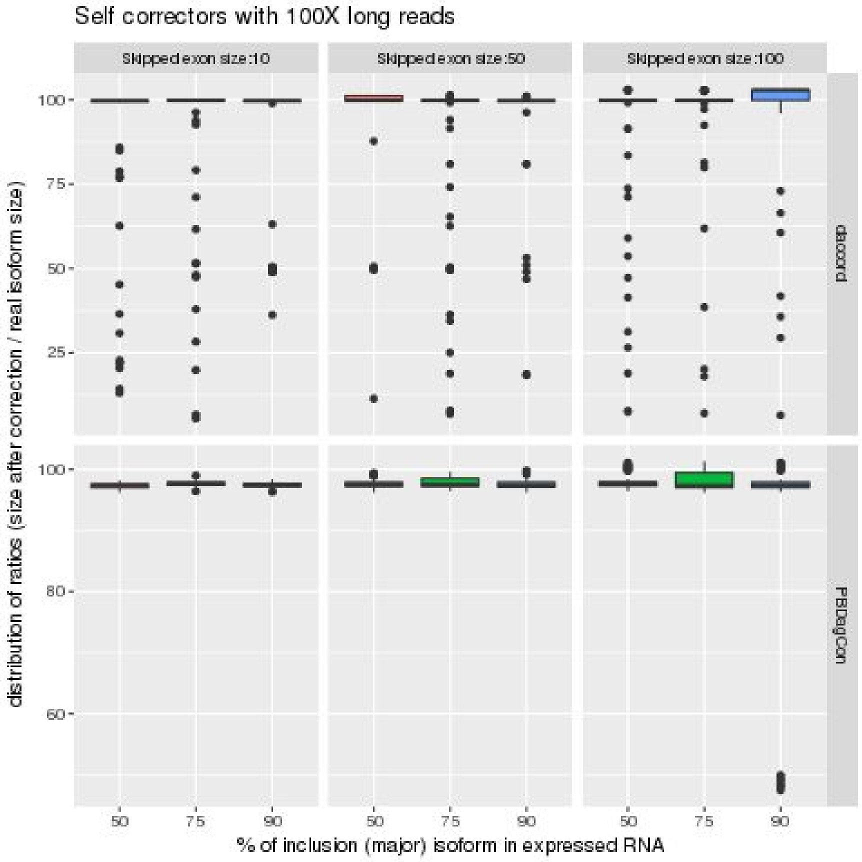


Figure S10. Counterpart of Figure S5 but with 100x coverage of long reads.

#### S9. Detailing incorrectly mapped splice sites

This plot shows the distribution of incorrect splice sites per tool according to 3 categories: “Incorrect SSs Near True SSs”, “Incorrect SSs with distance Multiple of 3” and “Incorrect SSs Far From True SSs”. “Incorrect SSs Near True SSs” is the number of incorrect Splice Sites near (at most 2 bps) away from the nearest true Splice Site. Such small errors could be due to splice alignment problems or small correction problems. Care must be taken when considering these small variations as true new splicing junctions. “Incorrect SSs with distance Multiple of 3” is the number of incorrect Splice Sites that are at a distance multiple of 3 of the nearest true Splice Site. A high value here might suggest many new splice junctions, but not always. For example, a new alternative acceptor site with a different ORF from the annotated ORF will cause the donor site to have a distance not multiple of 3 from the nearest true Splice Site. “Incorrect SSs Far From True SSs” is the number of incorrect Splice Sites that are not near (more than 2 bps) away from the nearest true Splice Site and are at a distance that is not multiple of 3 of the nearest true Splice Site.

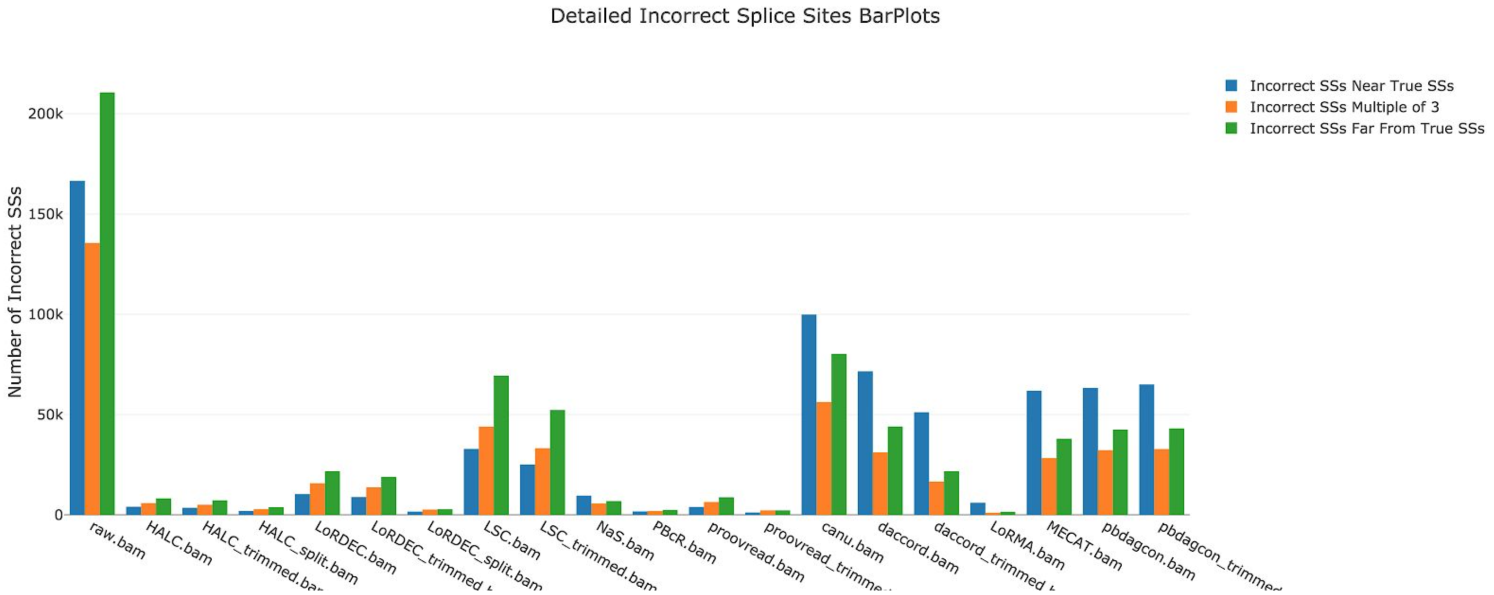


Figure S11. The distribution of incorrect splice sites per tool according to 3 categories: “Incorrect SSs Near True SSs”, “Incorrect SSs with distance Multiple of 3” and “Incorrect SSs Far From True SSs”.

The next plot shows the uncategorized distribution of incorrect splice sites per tool.

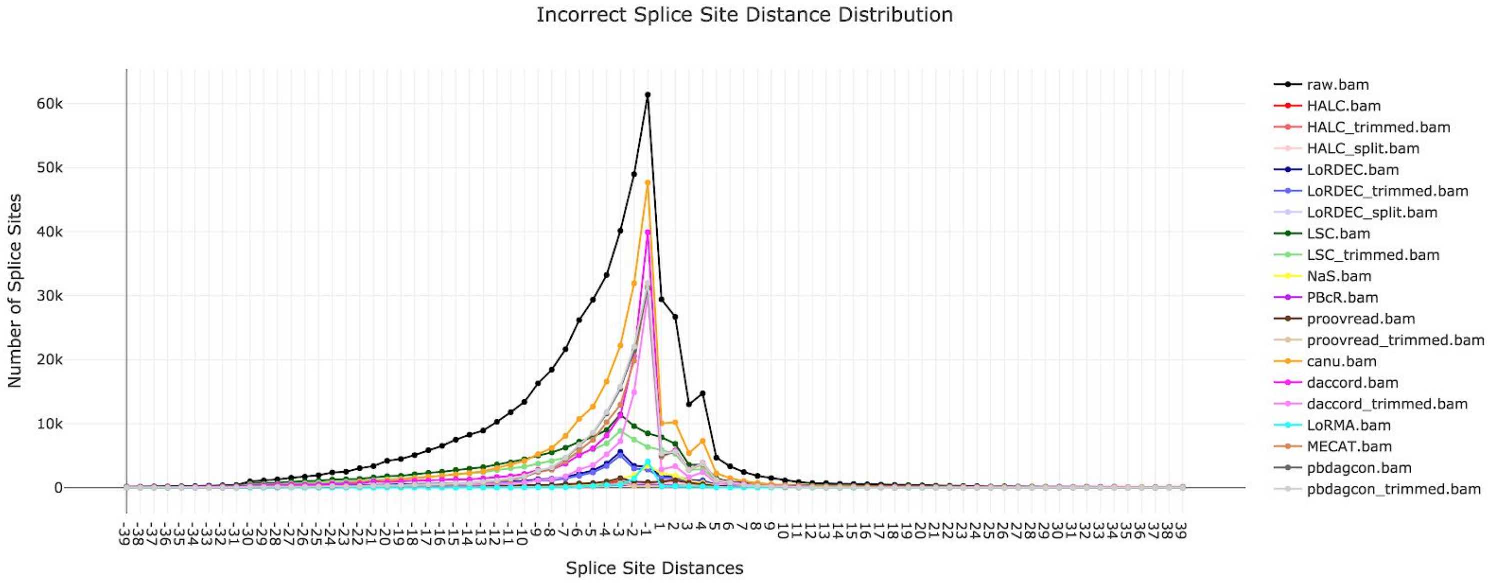


Figure S12. The uncategorized distribution of incorrect splice sites per tool according to 3 categories: “Incorrect SSs Near True SSs”, “Incorrect SSs with distance Multiple of 3” and “Incorrect SSs Far From True SSs”.

#### S10. Error-correction affects reads length and long reads connectivity

An interesting question is whether error correctors keep the long range information given by the long reads. One of the main advantages of this new technology in the RNA context, for example, is being able to sequence entire transcripts, unraveling the complete isoform structure and enabling distant exon coupling, which could be impossible, arguable or complicated with second generation sequencing. In some applications, it might not be worth to lower the error rate at the cost of losing connectivity, so in this section we investigate in more details if error correction tools decrease the reads' length and long reads connectivity.

A succinct answer to this question can be retrieved with the "mean length" metric in Table 2 of the main text. Inspired by AlignQC plots, we plotted in Figure S13 the amount of reads in different length intervals for each tool.

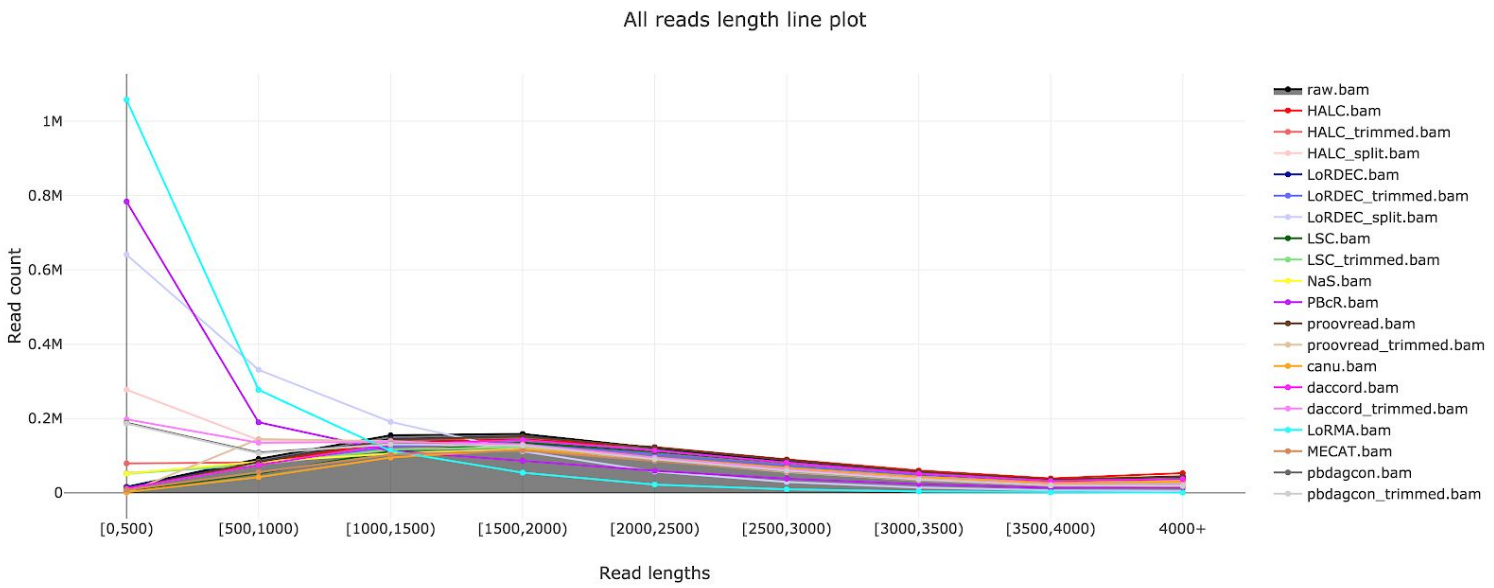


Figure S13. The amount of reads in different length intervals for each tool. In the x-axis we have the read length intervals, and in the y-axis we have the read count. The area between the points in the raw.bam line and the y=0 line was filled - points way above this filled area suggest that we have way too many reads in that interval.

We can see in Figure S13 that PBcR(s), LoRMA(s) and LoRDEC(s) are the correctors that tend to reduce the most the length of the long reads. They present an excessively high amount of reads with length [0, 500) and [500, 1000), while the number of reads longer than 1500bps decreases in comparison to the raw reads' length. Although these tools present high proportions of mapped reads, mapped bases, and low error rate after correction in their categories in Table 2 of the main text, they split long reads into well corrected subreads, potentially crippling downstream analysis that might need the long range information provided by the raw dataset. This is also evidenced by the number of identified exons per read in each tool (Figure S14) and by the number of reads with full and partial matches with respect to its mapped transcript in each tool (Figure S15).

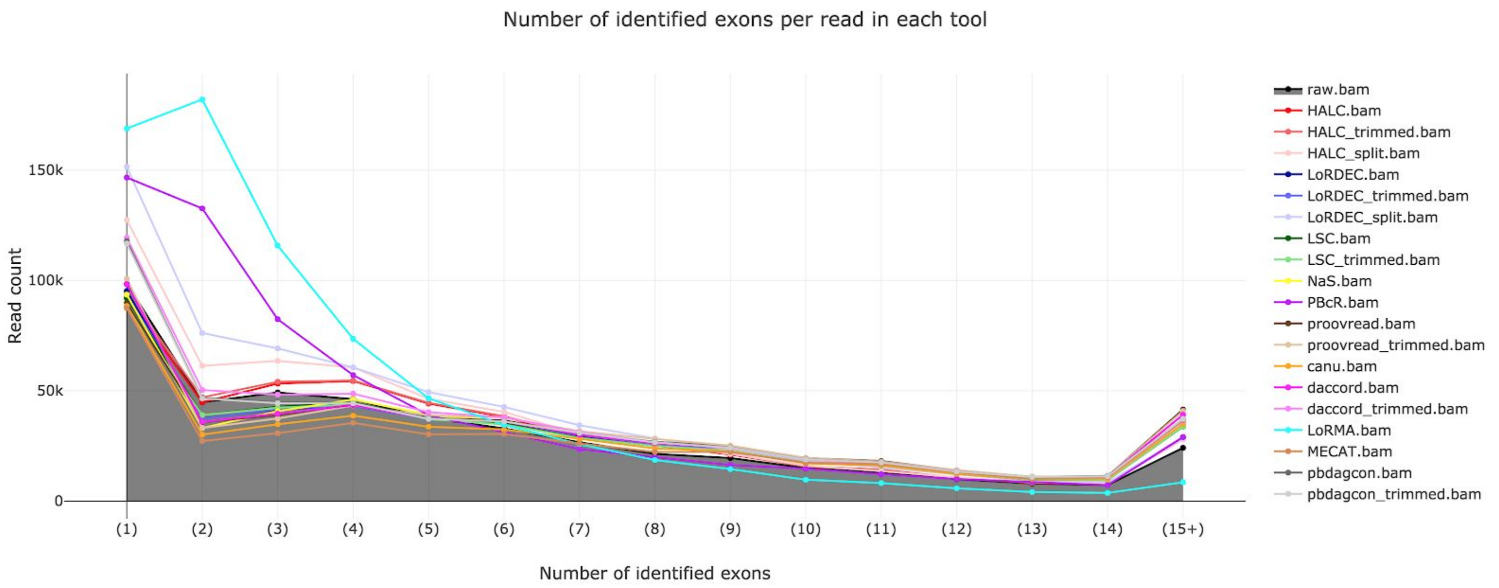


Figure S14. Number of identified exons per read in each tool.

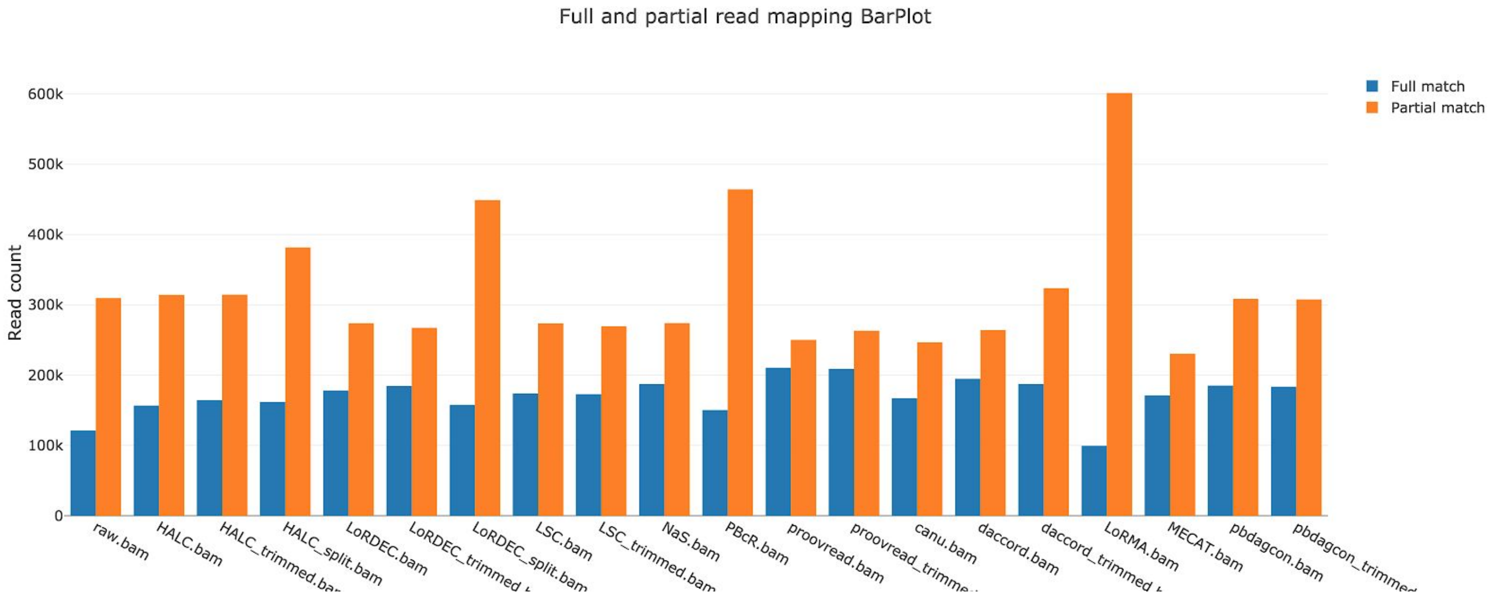


Figure S15. Number of reads with full and partial matches with respect to its mapped transcript in each tool.

### S11. Distribution of the difference of relative expression in the transcripts

This box and whisker plot represents the distribution of the difference of relative expression in the transcripts. It was done by taking, for each tool, each transcript that is expressed either in the raw reads or in the corrected reads, and computing the absolute difference between the relative expression of the transcript before and after correction. The relative coverage or expression of a transcript is the number of reads mapping to the transcript divided by the number of reads mapping to its gene.

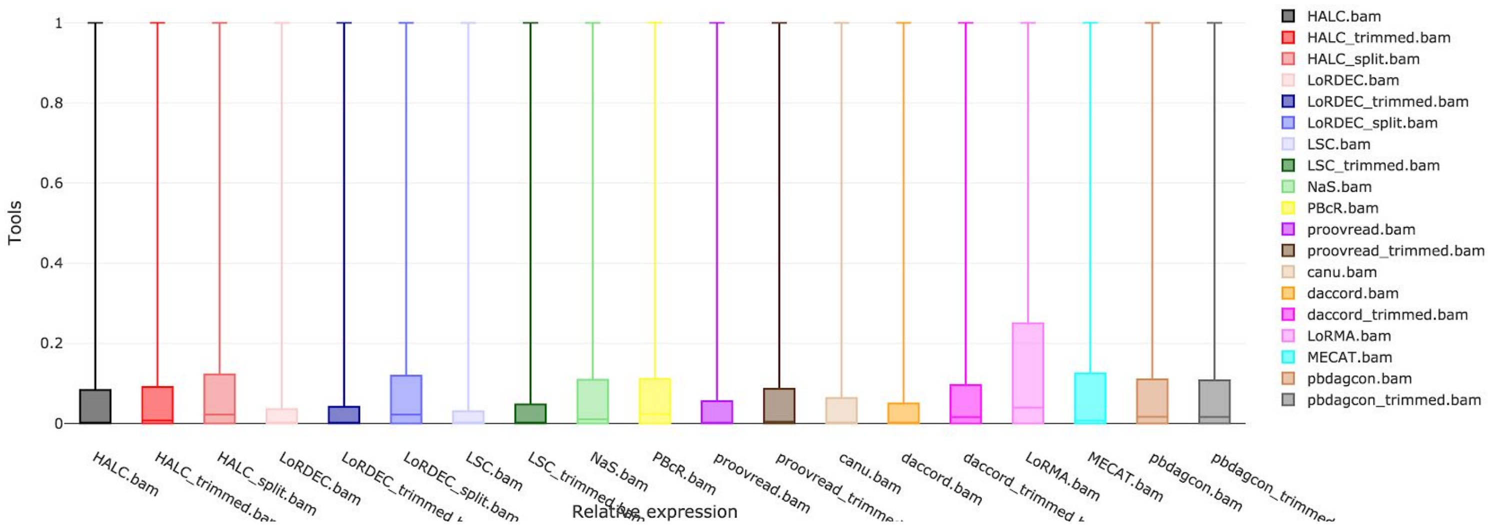


Figure S16. The distribution of the difference of relative expression in the transcripts.
